# A desert green alga that thrives at extreme high-light intensities using a unique photoin-hibition protection mechanism

**DOI:** 10.1101/2022.08.14.503881

**Authors:** Guy Levin, Michael Yasmin, Marc C. Simanowitz, Ayala Meir, Yaakov Tadmor, Joseph Hirschberg, Noam Adir, Gadi Schuster

## Abstract

While light is the driving force of photosynthesis, excessive light can be harmful. Photoinhibition, or light-induced photo-damage, is one of the key processes limiting photosynthesis. When the absorbed light exceeds the amount that can be dissipated by photosynthetic electron flow and other processes, damaging radicals are formed that mostly inactivate photosystem II (PSII). A well-defined mechanism that protects the photosynthetic apparatus from photoinhibition has been described in the model green alga *Chlamydomonas reinhardtii* and plants. *Chlorella oha-dii* is a green micro-alga, isolated from biological desert soil crusts, that thrives under extreme high light (HL) in which other organisms do not survive. Here, we show that this alga evolved unique protection mechanisms distinct from those of *C. reinhardtii* and plants. When grown under extreme HL, significant structural changes were noted in the *C. ohadii* thylakoids, including a drastic reduction in the antennae and the formation of stripped core PSII, lacking its outer and inner antennae. This is accompanied by a massive accumulation of protective carotenoids and proteins that scavenge harmful radicals. At the same time, several elements central to photoinhibition protection in *C. reinhardtii*, such as psbS, the stress-related light harvesting complex, PSII protein phosphorylation and state-transitions are entirely absent or were barely detected in *C. ohadii*. Taken together, a unique photoinhibition protection mechanism evolved in *C. ohadii*, enabling the species to thrive under extreme-light intensities where other photosynthetic organisms fail to survive.

## Introduction

Photosynthesis is the process by which plants, algae and cyanobacteria absorb and convert sunlight to chemical energy, which is stored in the chemical bonds of glucose newly formed via carbon fixation in the Calvin-Benson-Bassham cycle. Light is essential for photosynthesis and its lack thereof is one of the most limiting growth factors in a variety of shaded habitats ^1^. At the same time, during exposure to excessive light intensities, the amount of absorbed energy may exceed the capacity to utilize it through photochemical quenching. To protect themselves from excess light, photosynthetic organisms employ various mechanisms to dissipate light energy via non-photochemical quenching (NPQ) pathways ^2–7^. Additional protection mechanisms assist photosynthetic organisms in minimizing the damage elicited by excess light, by either reducing energy uptake by shrinking the photosynthetic antenna cross-section ^8–12^, or by rapidly repairing the compromised photosynthetic apparatus ^13,14^. Once the absorbed light intensity exceeds the threshold that can be safely handled via photochemical and non-photochemical quenching mechanisms, excited chlorophylls form triplet states, which transfer the excitation energy to oxygen to produce highly reactive singlet oxygen (^1^O_2_). ^1^O_2_ and other generated reactive oxygen species (ROS) can still be scavenged by carotenes and tocopherols as a last line of defense before they react with the D1 protein of the photosystem II (PSII) reaction center, damaging it and effectively causing photoinhibition (PI). To overcome PI, the damaged D1 must be removed, degraded, and replaced by a newly synthesized, active D1 ^13,15,16^.

*Chlorella ohadii* is a unique green alga closely related to the *Chlorella sorokiniana* species, that was isolated from biological soil crusts (BSC) in the Negev desert in Israel ^17,18^. The desert BSC is a harsh habitat characterized by subfreezing temperatures in the winter and scorching temperatures in the summer. Shade is scarce in this dry and arid landscape and an alga inhabiting this ecosystem must be well-adapted to the extremely intense light of the bright summer days, to avoid PI ^19,20^. Previous works uncovered several such adaptations, including the fastest generation of a photosynthetic eukaryote ^17 21^, a possible protective PSII-cyclic electron flow and multiphasic growth governed by metabolic shifts ^9,21–24^. We have shown how *C. ohadii* takes several known PI resistance mechanisms to the extreme. More specifically, when grown at extreme light of 2000 µE m^-2^s^-1^, *C. ohadii* reduces the PSII antenna, accumulates massive amounts of carotenoids and employs a very rapid PSII repair cycle^25^. Nevertheless, it is apparent that additional properties enable the maintained maximal growth rate and photosynthetic activity under such light conditions.

Here, we analyzed the unique *C. ohadii* resistance to extreme high light intensities. Interestingly, *C. ohadii* lacks some of the key players of the known PI protection mechanisms described in the model alga *C. reinhardtii* and plants. These include the light harvesting complex stress-related (LHCSR), psbS, state-transitions and thylakoid protein phosphorylation. In exchange, massive accumulation of a carotene biosynthesis related protein (CBR), an LHC-like protein that offers alternate routes for energy dissipation, occurred in high-light-grown cells. These unique PI resistance mechanisms of *C. ohadii* which enable it to thrive in light intensities that are fatal to most photosynthetic organisms, and how these mechanisms are distinguished from those of the model alga *C. reinhardtii*, are discussed.

## Results

### *C. ohadii* thrives at high-light intensities where other photosynthetic organisms are bleached by photoinhibition

*C. ohadii* habitat is the biological soil crusts (BSC) of the Israeli desert, which is exposed to very high-light (HL) intensities during the summer ^18^. Accordingly, this alga shows an unaffected growth rate even under very intense HL ^9,17,21–23,25^. In order to examine whether this species can grow at a light intensity where other organisms lose viability due to photoinhibition, log-phase *C. ohadii*, its closest relative species *C. sorokiniana*, as well as the other green algae, including the often used model organism *Chlamydomonas reinhardtii* and the cyanobacterium *Synechococcus elongatus* sp. PCC 7942, were analyzed for their ability to grow under HL. At a light intensity of 2500 μE m^-2^s^-1^, regardless of all other growth parameters, only *C. ohadii* and *C. sorokiniana* grew in cultures (Figs 1 and S1). For *C. sorokiniana* the results were not conclusive as in some experiments they did photo-bleach but on others they grew similar to *C. ohadii* (Figs 1 and S1). The other examined strains were consistently photoinhibited and bleached. These results indicated that *C. ohadii* and *C. sorokiniana* are more resistant to PI, an observation which prompted a search for unique protection properties that evolved in these species.

**Figure 1.**
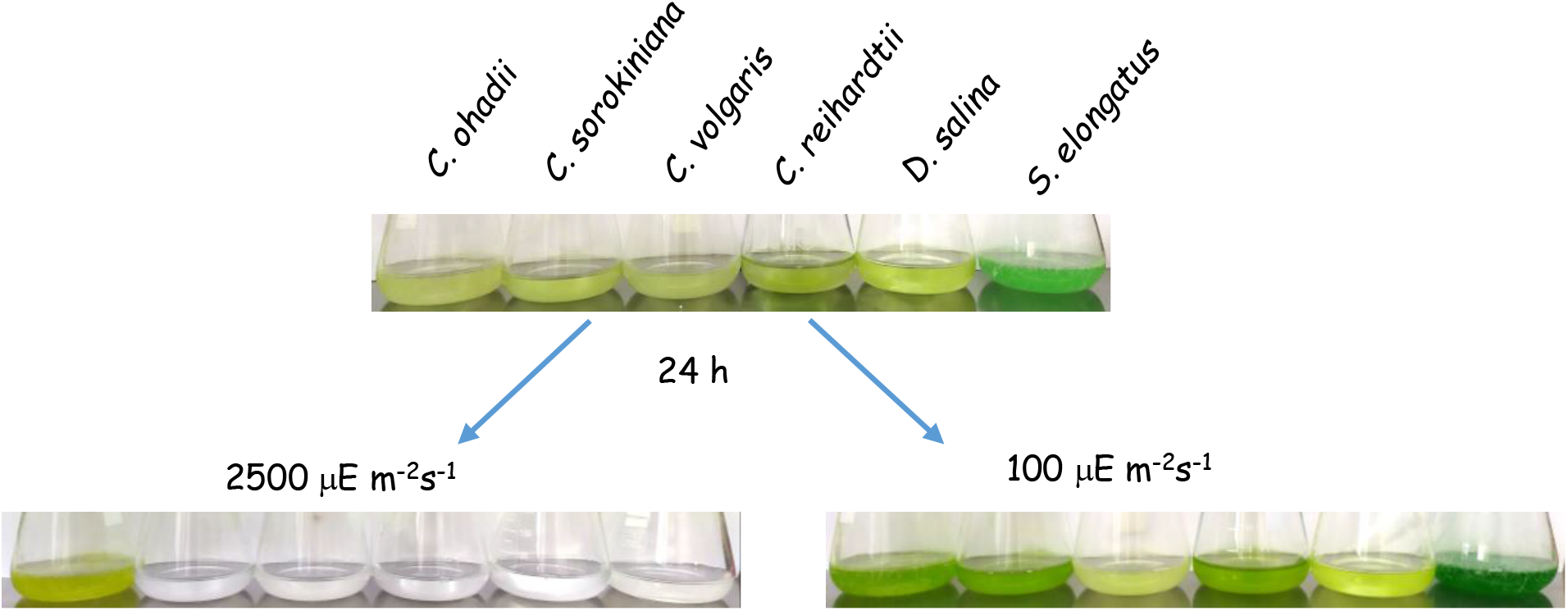
*C. ohadii* thrives at high light intensities where other photosynthetic organisms are lose viability due to photoinhibition. Log phase grown cultures at OD_750_=0.14 (upper panel) of the green algae *Chlorella ohadii, Chlorella sorokiniana, Chlorella vulgaris, Dunaliella salina* and the cyanobacteria *Synechococcus elongatus* were grown for 24 h at the light intensities of 2500 µE m^-2^s^-1^ (left lower panel) or 100 µE m^-2^s^-1^ (right lower panel). The results of *C. sorokiniana*, which is a very similar species to *C. ohadii*, where not significant and in about half of the experiments it grew similar to *C. ohadii*.

### *C. ohadii* displays limited state transitions and no phosphorylation of PSII proteins

In our previous work, we compared two *C. ohadii* populations that were acclimated to low or high light intensities by growing the cells either in low light (LL) or high light (HL) of 100 and 2000 μE m^-2^s^-1^, respectively ^25^. Both the LL and HL cells were highly active in the linear photosynthetic electron transfer rate and both displayed similarly rapid growth rates, with multiplication every 4 h. However, the LL and HL cells differed significantly in their photosynthetic properties, carotenoids contents and PSII structure ^25^. Since a major mechanism governing the balance between the light absorbed between the two photosystems and PI protection is the state transitions, which is mediated by the phosphorylation/dephosphorylation of PSII and light-harvesting complex II (LHCII) proteins, we analyzed this process in *C. ohadii*.

State transitions is a well-studied process involved in balancing energy uptake between PSI and PSII in response to frequent changes in light conditions ^2,12,26^. In state I (STI), PSI is preferably photo-activated, for example by enriched far-red light. The antenna of PSII is then enlarged by binding more LHCII proteins, resulting in a balanced and equal linear electron flow through the two photosystems. The transition to state II (STII), when PSII is preferably excited by, for example, an enriched red-light, is mediated by the phosphorylation of LHCII subunits and their shuttling from PSII-LHCII to PSI-LHCI, forming a PSI-LHCI-LHCII super-complex ^27–29^. This process can be reversed via the dephosphorylation of the LHCII subunits. In this manner, state transitions enable optimal distribution of the incoming flux of light energy between the two photosystems and significantly contributes to PSII protection by diverting excess energy to the more robust PSI. In *C. reinhardtii*, it has been postulated that during STII, a large fraction of the LHCII subunits are shuttled to PSI ^30,31^. To investigate whether *C. ohadii* employs this mechanism, state transitions were induced in LL and HL cells, as well as in *C. reinhardtii* as a positive control. Surprisingly, compared to *C. reinhardtii*, very limited STT was observed in LL *C. ohadii*, as evidenced by the small shift in the ratio between the fluorescence peaks of PSII/PSI while no difference was observed in HL cells (Fig 2a). More specifically, while for *C. reinhardtii*, the PSII/PSI ratio was 0.7 (STI) and 2.1 (STII), for LL *C. ohadii*, this ratio was 2.0 (STI) and 2.5 (STII). Similar results were previously obtained with LL-grown *C. sorokinianna* ^32^. However, the total lack of state transitions in HL *C. ohadii* is a unique property that has only been reported for *Chlamydomonas* species isolated in Antarctica ^33^.

**Figure 2.**
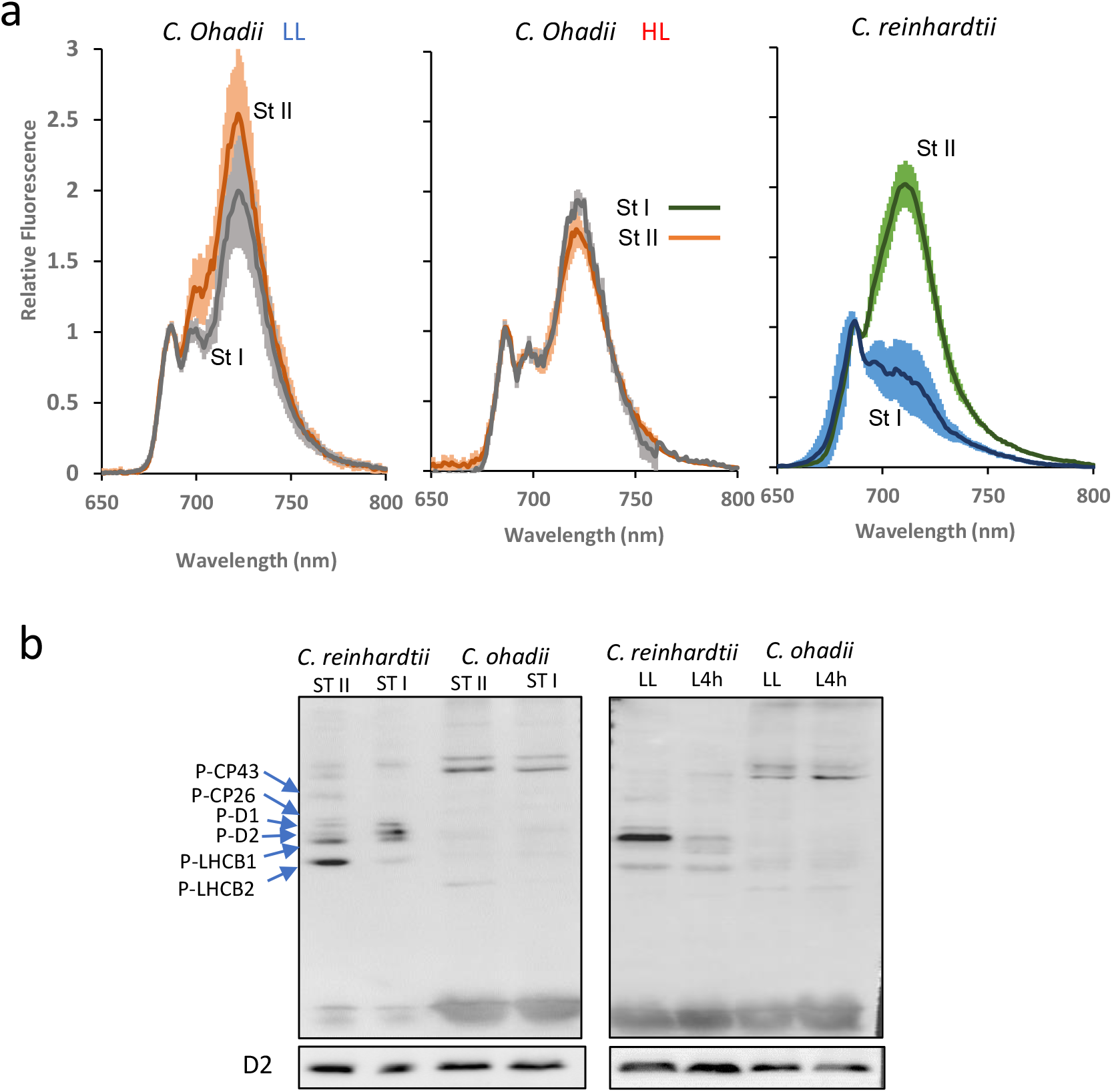
State transitions and phosphorylation of PSII proteins are very limited in *C. ohadii*. **(a)** State I (st I) and state II (St II) were induced by DCMU and FCCP treatments, respectively, and followed by 77k fluorescence emission spectra analysis of LL (left) and HL (center) cells of *C. ohadii*, as well as *C. reinhardtii* (right) as a positive control. Excitation was at 430 nm. The fluorescence emission spectra were normalized to the 687 nm peak of PSII. Data are means of at least three biological replicates with the standard deviation presented. **(b)** Detection the phosphorylated thylakoid proteins using phosphothreonine/tyrosine antibody. *C. ohadii* thylakoids of LL and HL cells adapted to ST II and ST I and those of *C. reinhardtii* (as a positive control) were analyzed (left). In addition, phosphorylated thylakoid proteins of *C. ohadii* and *C. reinhardtii* cells that were either grown in LL or transferred to HL for 4 h were detected (right). Immuno-detection of the PSII D2 is presented at the bottom to show equal thylakoid proteins loading. The identification of the PSII phosphoproteins were done as shown in Fig S2 and is indicated to the left.

State transitions are driven by phosphorylation/dephosphorylation of the LHCII and PSII proteins ^12^. In STII conditions, LHCII proteins are phosphorylated, disconnected from PSII and move to partially serving PSI. In light of the limited or lack of state transitions, the phosphorylation status of the thylakoid PSII proteins were next characterized (Figs 2b and S3). Homologs of the stt7 and stl1 corresponding protein kinases of *C. reinhardtii* were found in the proteome of *C. ohadii* thylakoids (Fig S2). Phosphorylated PSII proteins and, in particular, the LHCII at STII, were observed in *C. reinhardtii*. However, in agreement with the very limited state transitions, no phosphorylation of PSII proteins, including LHCII, was detected in *C. ohadii*, regardless of whether the cells were in STI or STII. Since absence of phospho-LHCII was unexpected, their phosphorylation status under another physiological condition, 4h after transfer from LL to HL, was assessed. In this case as well, no phosphorylation of the LHCII subunits or any other PSII proteins was detected (Fig 2b). Two proteins of an approximate molecular weight of 50 kDa reacted with the phosphothreonine antibodies in both HL and LL *C. ohadii* samples. These may either be thylakoids protein that are exclusively present or phosphorylated in *C. ohadii*, as compared to *C. reinhardtii*, or a contamination of the thylakoid preparation with soluble phosphoproteins. Together, these results indicated a limited role of state transitions in PI protection of *C. ohadii*. While the lack of state transitions in HL cells could be explained by the large reduction of LHCII, its limited activity in LL cells was unexpected, as it is believed to significantly contribute to protection from PI in *C. reinhardtii* and plants ^2^. The lack of PSII protein phosphorylation seems to be a unique characteristic of *C. ohadii*.

### A core PSII, in which both the major and minor antenna are removed, replaces the PSII-LHCII complex in HL cells

Previous physiological, spectroscopic and biochemical analyses indicated that there is a reduction of the PSII antenna of HL cells ^25^. Fractionation of the PSII complex of LL and HL cells identified four chlorophyll-protein complexes, in addition to unbound chlorophyll and carotenoids, with marked enrichment of carotenoids in the HL samples (Fig 3a). Characterization of the four complexes detected the four chlorophyll-containing complexes of *C. ohadii*, PSII-LHCII, PSI-LHCI, core PSII and detached LHCII (Fig 3) ^26^. The detached LHCII could be either LHCII that was not associated with PSII or an LHCII that was disconnected as a result of the sample preparation process.

**Figure 3.**
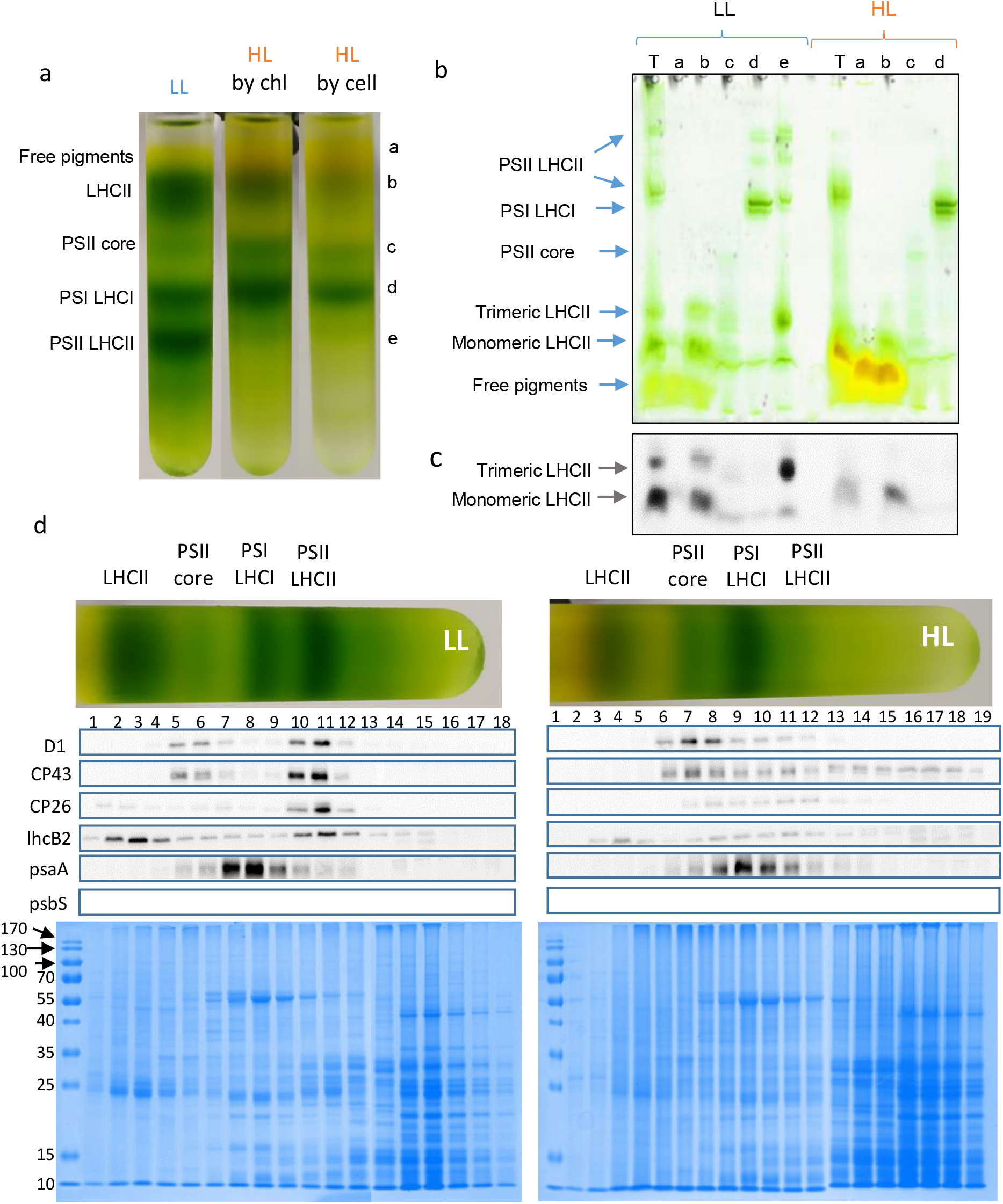
High light grown cells contain a core PSII that lacks the light harvesting antennae. **(a)** The chlorophyll-protein complexes were extracted from thylakoids of low light (LL) and high light (HL) grown cells and separated by sucrose gradient centrifugation. The identities of the chlorophyll containing complexes are indicated on the left. Two gradients of the HL sample are shown. One gradient was loaded with an equal chlorophyll content in order to observe the complexes (by chl). The second was loaded with half amount of chlorophyll as to the LL sample in order to present the amount of the complexes per cell, since the HL cells contain half the amount of chlorophyll of LL cells (by cell). **(b)** The complexes were analyzed by non-denaturing native gels. The identity of each fraction is indicated at the top of the gels by the letters as indicated on the side of the sucrose gradient in panel a. T-thylakoids. **(c)** A picture of the fluorescence obtained from the gels when illuminated with red light. Only the lower part of the gel, containing the LHCII complexes is shown as the upper part did not show any fluorescence emission. **(d)** Each gradient was separated into fractions and the protein contents were analyzed by immunoblots, using specific antibodies, as well as by Coomassie blue staining of the SDS-PAGE. The size of the molecular weight markers in kilodaltons is indicated to the left.

The most apparent difference between the LL and HL thylakoids was the massive reduction in the amount of PSII-LHCII and the corresponding accumulation of the smaller core PSII complex in HL (Fig 3a-d). Interestingly, the HL core PSII was found to be stripped of both the major and the minor antennas, as revealed by immunoblotting with LHCb2 and CP26 antibodies, respectively (Fig 3d). These results revealed that a core PSII stripped of both outer and inner antennas, replaced the holo-PSII-LHCII complex in HL cells in order to confer PI protection and enable growth at HL intensities. Since the PSII-LHCII was the major form in LL, it was likely converted to the core PSII during the acclimation to HL. In addition, the amount of detached LHCII was significantly reduced in HL LHCII fraction, further strengthening the notion that a dramatic structural change took place in response to prolonged HL. It is reasonable to suggest that this change in the PSII antenna evolved in *C. ohadii* in order to enable better protection against PI during growth under extreme HL. While modulating the antenna size in response to different light intensities is a well-documented in algae and plants, the formation of a core-PSII that lacks the entire LHCII is unique to *C. ohadii* and its close relative *C. sorokiniana* ^34^.

The changes in the protein content of the PSII complexes were further analyzed by mass spectrometry (MS). Intensity based absolute quantification (iBAQ) ^35,36^ was used to determine the abundance of each identified protein and relative-iBAQ (riBAQ) was then calculated to determine the relative abundance (percentage of molarity, see Methods) (Figs 4a, S4 and Table S1). When comparing the dominant PSII form in LL (PSII-LHCII) and the dominant form in HL (core PSII), a dramatic reduction in the PSII antenna was noted in HL. Specifically, the relative abundance of lhcbm2, a subunit of the LHCII major antenna, in the core PSII fraction was 0.4%, 11-fold lower than its abundance in the PSII-LHCII complex of LL cells (4.4%). In parallel, the relative abundance of lhcbm4, another LHC major antenna subunit, was 0.2% (HL) vs. 3.8% (LL), i.e., a 19-fold difference between the two samples (Fig 4a). In agreement with the protein levels reported above (Fig. 3d), the marked reduction of LHCII in HL was not limited to the major antenna. The minor antenna subunits were also reduced in HL core-PSII (relative abundance of 0.5% (CP26) and 0.2% (CP29)) compared to LL PSII-LHCII (4.0% and 2.2%, respectively), yielding an 8-fold and 11-fold decrease, respectively. Most of the LHCII subunits were present in the detached LHCII fraction and, although to a lesser extent, the reduction of the subunits in HL cells was also apparent in this fraction. More specifically, relative abundance of the subunits in the detached LHCII fraction was 6.4% (lhcbm2) and 3.5% (lhcbm4) in HL cells compared to 10.1% and 15.4% in LL cells, yielding a 1.6-fold and 4.4-fold decrease, respectively (Fig 4a). The total degree of reduction in the amount of each antenna protein in HL is shown in the riBAQ values of total thylakoids proteins (Fig. 4a right). It is reasonable to assume that an additional increase in the light intensity would result in further reduction of both the minor and major LHCII antennae.

**Figure 4.**
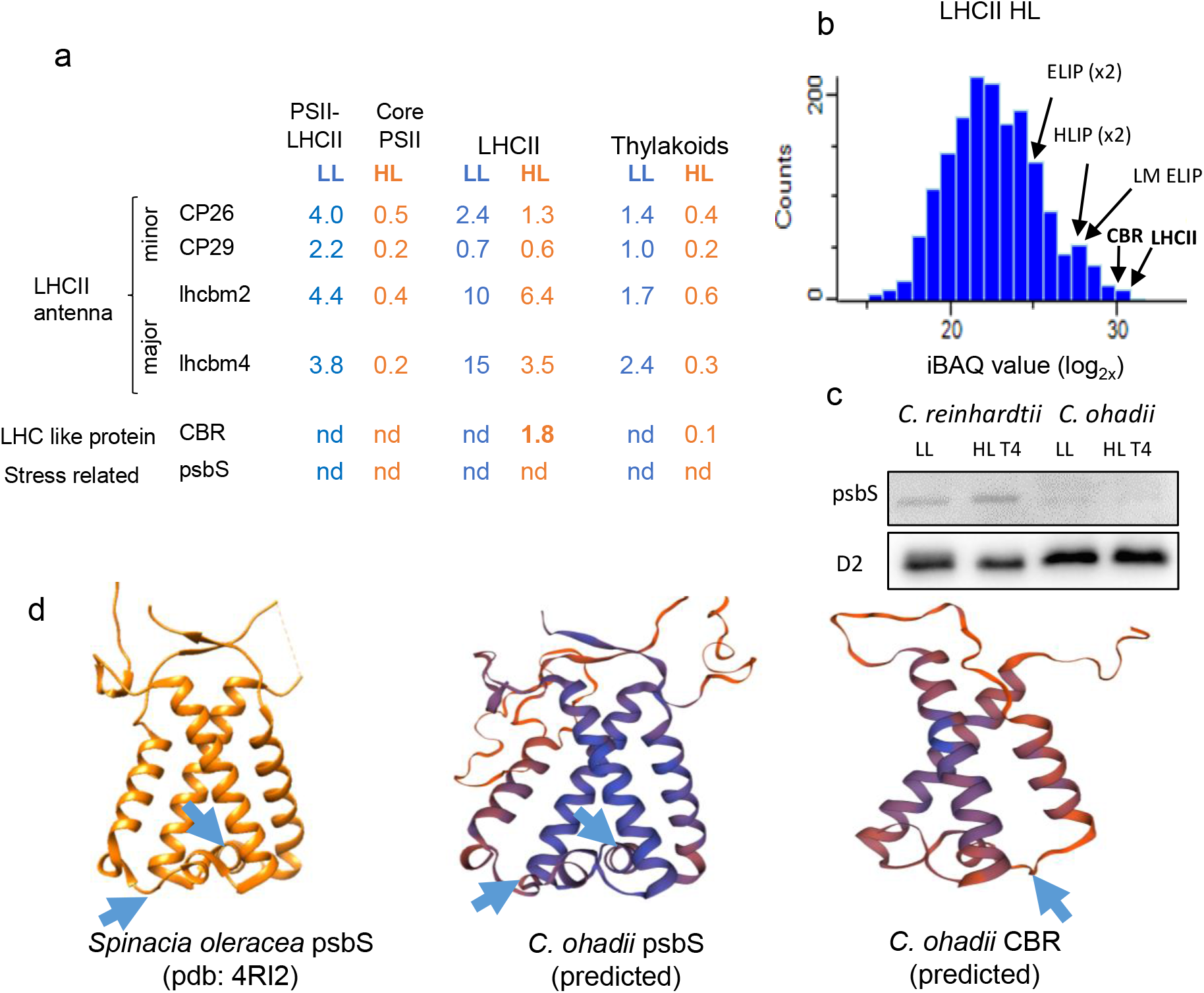
Both the minor and major antenna are reduced in HL and there is a large accumulation of CBR. **(a)** The relative molar abundance of several proteins in the purified complexes and thylakoids of LL and HL cells, as calculated from the proteomics analysis using relative iBAQ. **(b)** Peptide distribution in LL and HL thylakoids. The X axis displays the LFQ intensity (log_2_ scale) and the Y axis the peptide counts. Data analysis was performed and figures were generated with Perseus software (1.6.2.1). CBR-Carotenoids Biosynthesis Related protein. ELIP, HLIP and LM-ELIP are Early Light Induced Proteins that were identified, as the CBR and the other proteins by homology to known proteins from plants and algae in the data bank. The results of the riBAQ analysis of these proteins is shown in Table S2. **(c)** psbS was not detected using specific antibodies. Thylakoids of LL and cells illuminated 4h with HL were analyzed for the present of psbS. *C. reinhardtii* thylakoids were tested as a positive control and the D2 as a loading control. **(d)** The structure of *C. ohadii* CBR (right), based on the predicted *C. ohadii* psbS, as drown using the known structure of spinach psbS (left and middle), predicts a structure of three cross membrane alpha helices, similar to LHCII proteins. The conserved glutamate residues that sense the pH change in the lumen are indicated by blue arrows.

### LHC-like proteins accumulate in HL thylakoids

Various LHC-like proteins such as the early light-induced protein (ELIP), were postulated to function in photoprotection by providing alternate routes for energy dissipation ^11,34,37^. Interestingly, the decrease in LHCII subunits in the detached LHCII fraction under HL was accompanied by an accumulation of LHC-like proteins. Six differentially expressed LHC-like proteins were detected, five of which accumulated in HL and one of which decreased (Table S2). Most apparent was the accumulation of the carotenoid biosynthesis-related (CBR) protein, which was not detected in any of LL samples, but was the sixth-most abundant protein in the HL LHCII fraction, with a relative abundance of 1.8%, only half of the amount of lhcbm4 and about an order of magnitude higher than other LHC-like proteins (Figs 4a, S4 and Table S2). The remarkable accumulation of CBR in the HL detached LHCII complex is clearly observed in the intensity histogram of the distribution and number of proteins in this fraction (Fig 4b). The CBR appeared in the most abundant group of proteins surpassed only by the LHCII subunits. Analysis of the thylakoid membranes reveals CBR accumulates to 1/12 compared to D2, suggesting 1 CBR protein per 12 PSII reaction centers (Table S2). Similar stoichiometry was reported for psbS in HL stressed *C. reinhardtii* ^38^. CBR was previously shown to bind the carotenoids lutein and zeaxanthin alongside chlorophyll *a* ^39–43^. CBR may bind some of the accumulated xanthophylls and chlorophylls of the degraded LHCII, dissipating energy from excited chlorophylls to carotenoids before ROS are formed, thereby protecting against PI (Figs 4 and S5-S7). Structural modelling of CBR showed its resemblance to LHCII, with three conserved transmembrane helices, and a very non-conserved area located at the non-membranal luminal side (Fig 4d). In addition, the structure of the transmembrane helices is highly homologous to that of three transmembrane helices of the stress-related protein psbS (see below) (Fig. 4d). Interestingly, structural modeling suggested a glutamate that was further revealed in BLAST analysis as one of the two conserved glutamates of psbS that senses the pH change of the lumen resulting in the activation of PI protection processes and is also present in CBR (Figs 4d and S7). In addition, similar to LHCSR, the CBR sequence carried the ability to bind chlorophyll (Fig S5). For a full list of PSII and LHC II proteins in the complex samples analyzed by MS, see Table S1. Together, these results revealed an efficient protection mechanism, which included the elimination of LHCII in HL cells and the formation of core PSII, alongside massive accumulation of CBR in the remaining LHCII. Similar to the conclusion reached regarding lack of state transitions, these observations suggest a PI protection mechanism that is significantly different from that of the model algae *C. reinhardtii*.

### *C. ohadii lacks a* gene ortholog to the stress-induced LHCSR, and psbS and PTOX proteins are barely detected or absent in *C. ohadii*

MS analysis found similar inner stoichiometry of PSII subunits in the core PSII and PSII-LHCII of HL and LL cells, respectively, suggesting no major structural changes in the reaction center PSII core on exposure to HL (Table S1). The analysis detected the PSII major intrinsic subunits (psbA, B, C and D), the extrinsic subunits of the oxygen-evolving complex (OEC, psbO, P, Q and R), the cytochrome b559 subunits (psbE and F) and the low molecular mass psbH (Table S1). In *C. reinhardtii*, LHCSR and psbS are two important proteins which compose the q_e_ component of NPQ, and play a role in sensing the acidification of the lumen under HL conditions ^2^. Interestingly, LHCSR orthologs were not detected in the *C. ohadii* and *C. sorokiniana* genomes ^17,25^. While two orthologs of psbS were previously found in the genomes of *C. ohadii* and *C. sorokiniana* and their transcripts detected ^23^, the current analyses failed to identify psbS protein in thylakoids or the complexes (Table S2, Figs 4a and 4c). Since previous works reported transient expression of psbS when algae strains were illuminated for several hours ^44,45^, an attempt was made to detect expression following 4h of HL. Marked accumulation of the protein was observed in *C. reinhardtii*, while very low levels or complete absence of the proteins were found for *C. ohadii* (Fig 4c). These observations imply that this highly PI-resistant organism developed an alternative mechanism to induce the q_e_ component of NPQ via lumen acidification sensing, which is a key process in photoprotection of *C. reinhardtii* and plants ^46,47^. As described above, CBR may be a candidate sensor of the acidification at HL. Finally, the gene encoding PTOX, an additional player in the PI protection processes, has been found in the genome of *C. ohadii*, however, its expression level was very low (Fig S2).

### Upon illumination, fewer superoxide radicals accumulate in HL cells

ROS form during illumination and promote PI ^2^. Scavengers such as carotenoids that reduce the amount of accumulated ROS, protect the system from PI. Quantitation of superoxide (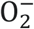) and hydrogen peroxide (H_2_O_2_) levels in thylakoids revealed 50% less 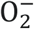 in HL cells as compared to LL cells, suggesting that either there is reduced production of 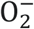 in HL cells and thylakoids or that there is a better scavenging of this radical, perhaps by the accumulating carotenoids (Figs 5a, S8 and Table S3). No difference was obtained in H_2_O_2_ levels between the two growth conditions.

**Figure 5:**
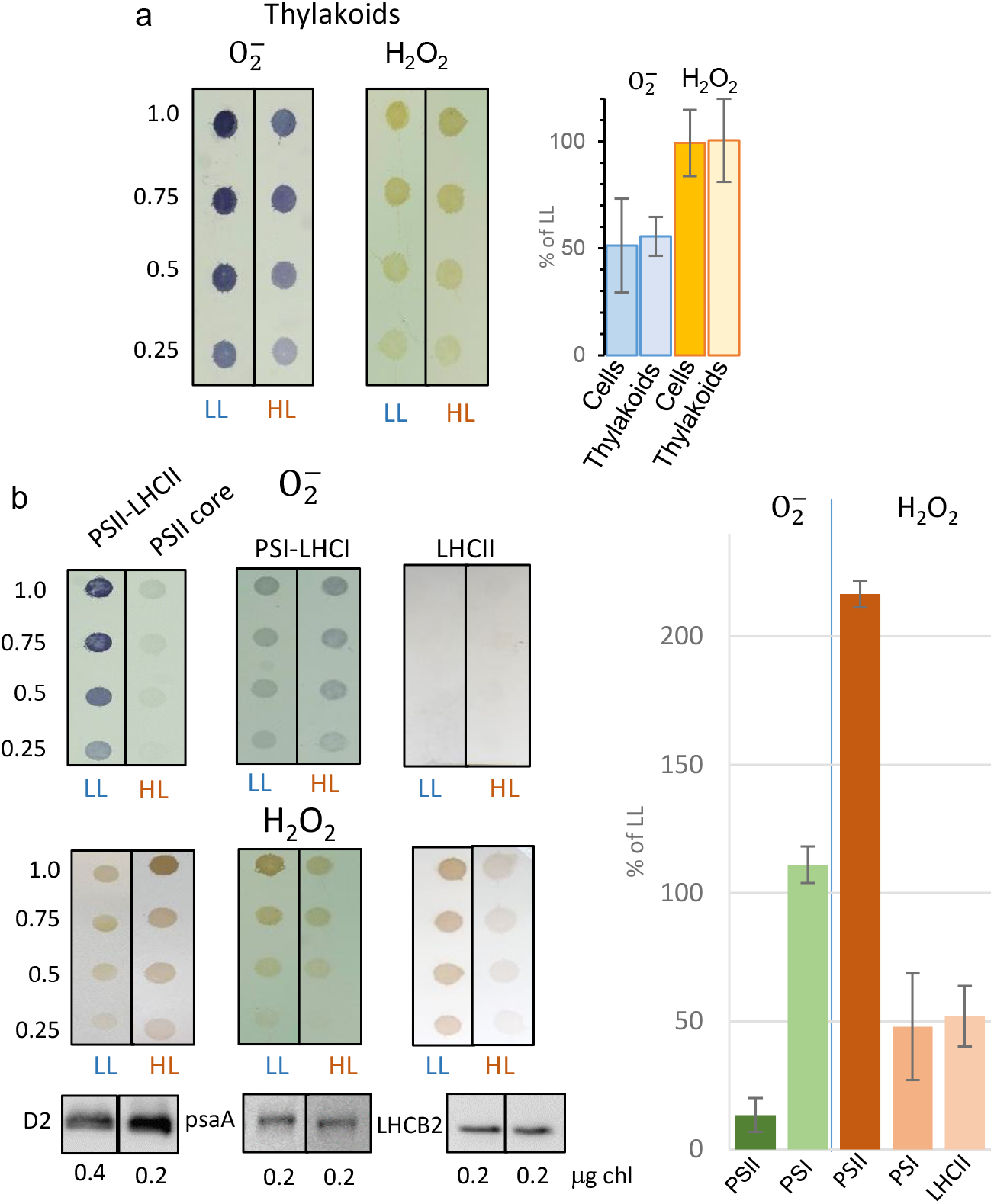
Different accumulation of ROS in LL and HL cells, thylakoids and the photo-synthetic complexes. Cells, thylakoids **(a)** and photosynthetic complexes **(b)** of LL and HL cells were spotted on PVDF membrane in a serial dilution as shown to the left and illuminated in the presence of NBT to detect the amount of superoxide radical (O_2-_), or DAB to observe hydrogen peroxide (H_2_O_2_). One repetition of at least three is presented. For the superoxide analysis the chlorophyll concentrations of the x1 dilution slots were 0.8 µg except the PSII core of HL which was 0.4 µg. For the H_2_O_2_ samples the chlorophyll concentration was 1.6 µg except the PSII core of HL which was 0.8 µg. Immunoblot using the corresponding protein is presented at the bottom to ensure equal loading. Quantitative analysis, presented on the right, was performed using the Image J software and included the standard deviation of at least three repetitions. The picture of the ROS analysis of cells is presented in Figure S8.

Analysis of the purified complexes detected a significant accumulation of 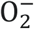 in the LL PSII-LHCII complex as compared to the HL PSII core, while the H_2_O_2_ accumulated more in HL (Fig 5b). The opposite trend was measured for PSI where similar levels of 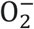 accumulated under both light conditions, but more H_2_O_2_ accumulated in LL. Surprisingly, no 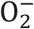 was detected in either LL or HL detached LHCs, while H_2_O_2_ levels were approximately double in the LL as compared to the HL LHC (Fig 5b). Together, these results revealed differences in the accumulation of two ROS upon illumination. The more modest accumulation of 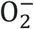 in HL cells, thylakoids and the PSII core may be related to the efficient PI protection mechanism that enables *C. ohadii* to thrive under HL intensity. As described above, a possible mechanism to reduce ROS in HL is the large accumulation of scavenging carotenoids and the CBR protein.

### The change in PSII structure is a long process

*C. ohadii* acclimation to HL involves several major changes in the thylakoids and photosynthetic properties. The major observed changes are the large quenching of the PSII fluorescence, the massive accumulation of protective carotenoids and the enormous reduction of the PSII antennae and the contaminant accumulation of the core PSII complex ^25^. It is important to analyze the temporal development of these three phenomena on transfer to HL. On one hand, the changes enabling rapid growth at HL are required to protect the cells from PI, yet, on the other hand, destruction and reconstruction of the PSII antennae on a daily basis is mostly cost ineffective and unlikely to take place each day during the summer. To examine the time-scale of the acclimation process, carotenoids accumulation and conversion of LHCII-PSII to core-PSII complexes were monitored after transfer to HL.

Extreme fluorescence quenching of PSII was noted within 10-20 min ^25^. The PSII-LHCII complex remained constant and there was no degradation of the PSII antennae during the initial 8 h, as evident by the stable chlorophyll *a/b* ratio and PSII-LHCII complex content (Fig 7a). However, during this same window, there was substantial accumulation of carotenoids (mainly lutein and zeaxanthin) in the cells, peaking after 4-5 h of HL (Fig 7b and Table S3). Thereafter, a gradual reduction in PSII-LHCII complex and chlorophyll *b* content was observed over the subsequent 15 h, until, eventually, no PSII-LHCII was detected (Fig 7a). The gradual disappearance of PSII-LHCII was accompanied by accumulation of the core-PSII complex, which became the major PSII. At 16-20 h following the transfer to HL, the carotenoids/chlorophyll ratio continued to increase due to the decrease in chlorophyll levels per cell, which was 50% lower in HL as compared to LL cells ^25^.

**Figure 7.**
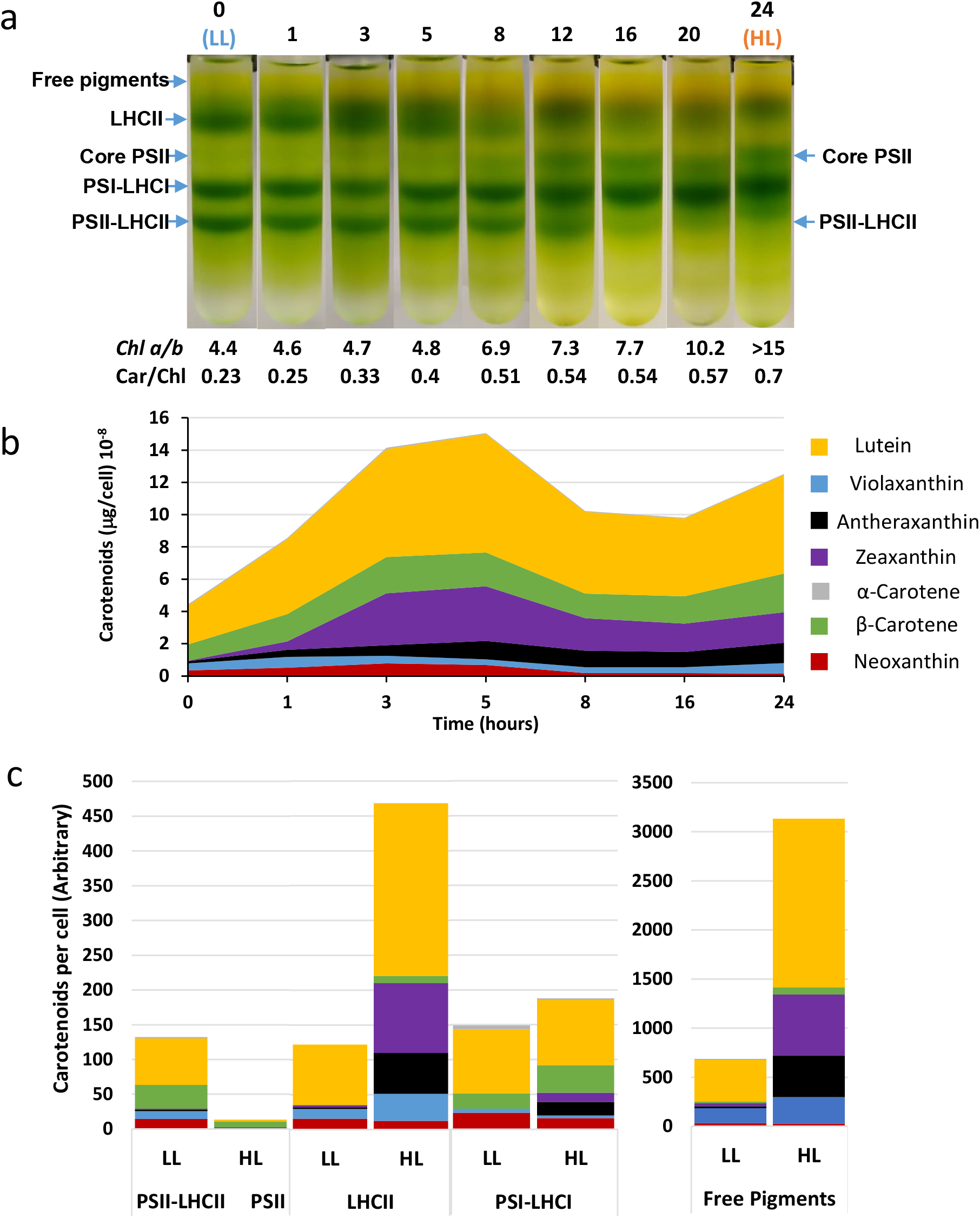
Changes in the photosynthetic complexes and the accumulation of carotenoids during the transition from LL to HL. Low light (LL) grown cells were transferred to high light (HL) and samples were withdrawn at the indicated times. **(a)** Thylakoids were obtained and the protein complexes were extracted and analyzed by sucrose density gradient separation. The chlorophylls *a/b* and the carotenoids/chlorophyll ratios are indicated. Note the disappearing of the PSII-LHCII and the appearance of the core PSII after 8 h. **(b)** Carotenoids were extracted from the thylakoids at each time point, analyzed and are presented as the amount per cell. **(c)** Changes in the carotenoids content of the photosynthetic complexes and the free pigment fractions of LL and HL cells. Note the different scale of the photosynthetic complexes and the free pigment fraction.

We suggest that these distinct acclimation phases evolved to counteract excess illumination on different time scales. While carotenoid accumulation peaked during a single bright day, the antenna reduction required much longer than the maximum daylight interval. Thus, while carotenoids accumulation and xanthophyll cycle activation can operate as a diurnal PI protection processes, the removal of the PSII antennae and formation of core-PSII is a seasonal rather than a daily adaptation mechanism. As the days grow longer towards the onset of the bright desert summer, the antenna reduction process initiates and is not reversed during the short summer nights.

### The protecting carotenoids accumulate in the LHCII and the phospholipids moiety of the thylakoids

Analysis of carotenoid content in the individual photosynthetic complexes of cells up to 24 h after transfer to HL conditions, identified a significant accumulation of carotenoids in the free pigment fractions, and to a lesser extent, in the detached LHCII fraction (Fig 7c and Table S3). In contrast, the detached LHCII fraction of HL cells contained increased levels of lutein (x6), zeaxanthin (x28) and antheraxanthin (x30), the three major PI-protecting carotenoids, as compared to LL. Regarding the marked accumulation of CBR protein in the detached LHCII fraction of HL cells (Fig 4), it is possible that in addition of occupying the binding sites in the remaining LHCII subunits, the accumulated carotenoids also bind CBR. There was very little carotenoid content in the core PSII complex of HL cells, aligning with the loss of antenna (Fig 7C and Table S3). In addition, the considerable accumulation of lutein (x4), zeaxanthin (x18) and antheraxanthin (x24) in the free pigment fraction of HL cells strongly suggests that they are present in the phospholipid moiety of the thylakoids not bound to proteins (Fig 7c, Table S3).

In HL, β-carotene was the key carotenoid to accumulate in the core PSII and the PSI-LHCI complexes (Fig 7c, Table S3). Yet, this protective carotene accumulated to a much lesser extent than the lutein and the other xanthophylls in the LHC complexes and the free pigment fraction. Unlike the enormous differences between the LL and HL PSII complexes, the difference between the LL and HL PSI-LHCI complexes were minimal, as was also shown before (Fig 7c, Table S3)^48^.

Together, these results demonstrated that the xanthophyll cycle is activated with the onset of HL, resulting in massive accumulations of the PI protective carotenoids in the LHCII, the phospholipids moiety and possibly in the accumulating CBR. The transformation of the LHCII-PSII to the core-PSII is delayed, starting after about 8 h of HL, and is completed only after 20 h of HL, suggesting it to be a seasonal rather than a diurnal change.

## Discussion

### The PI protection mechanism of *C. ohadii* is different from that of *C. reinhardtii* and plants

Living in the harsh habitat of the dessert sand crust, *C. ohadii* was compelled to develop an efficient PI protection mechanism. Therefore, its photosynthetic properties have been intensively studied in search for new PI protection mechanisms that evolved in this organism, and for their potential implementation in agriculturally valuable plants and algae. These studies revealed that *C. ohadii* is the fastest growing photosynthetic eukaryote, with a multiplication rate, even at HL intensities, of 1.5-4 h, and exhibits unique changes in its metabolic profile ^9,17,21–23,25,48^. In addition, under HL, its photosynthetic activities were modified, partly in PSI ^48^, but mostly in PSII ^9,25^. In our previous work, we found that two central processes of PI protection, i.e., the activation of the xanthophyll cycle and the rapid repair cycle of PSII, are most efficient in *C. ohadii* ^25^. In addition, the PI protection mechanisms differed from those described so far in the model alga *C. reinhardtii*, as manifested by major structural changes in the PSII structure, enormous accumulation of protective carotenoids and marked quenching of PSII fluorescence. Therefore, this work concentrated on the differences in the PI protection of *C. ohadii* and the model alga *C. reinhardtii* (Table 1).

**Table 1.**
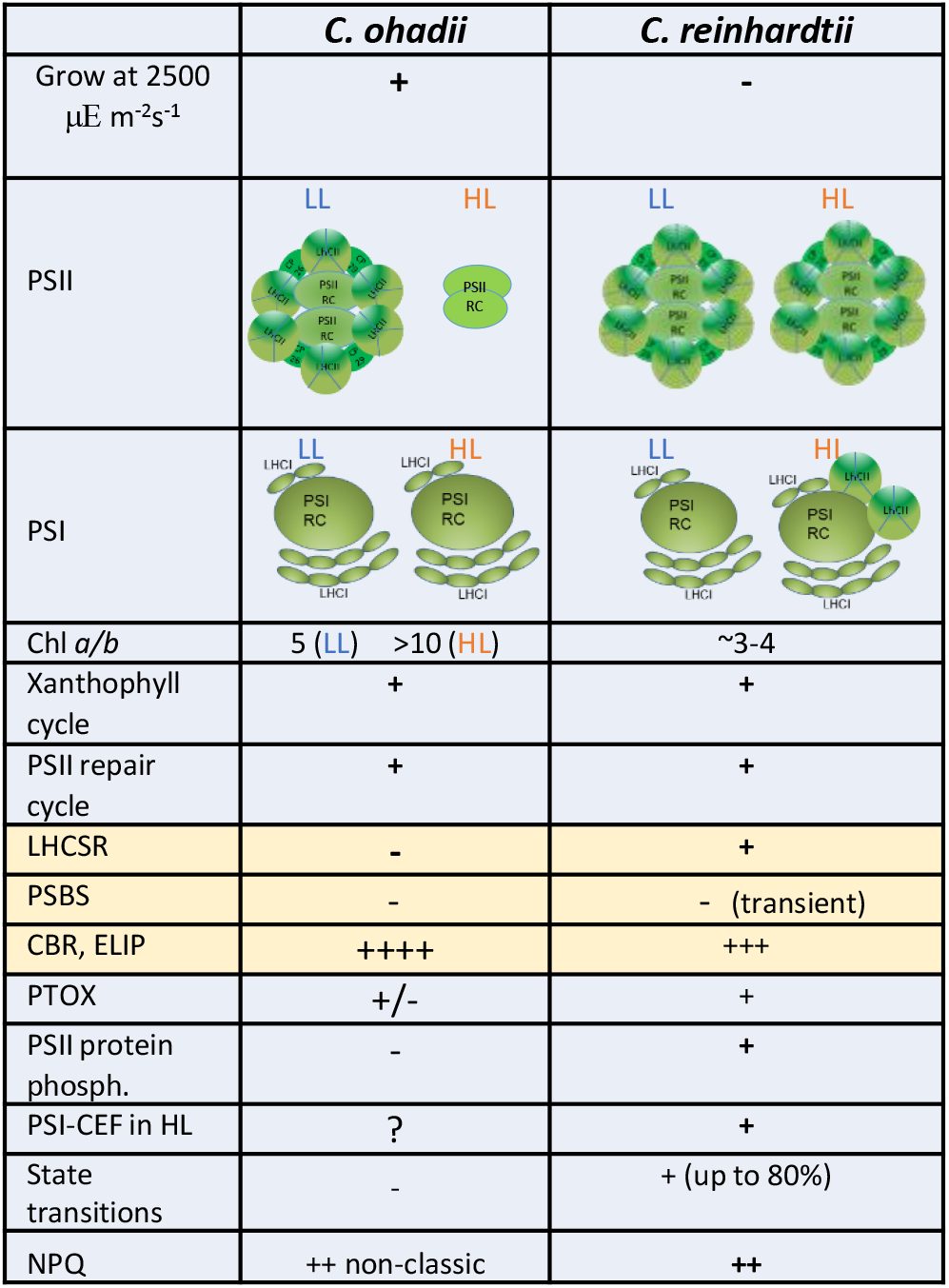
The successful PI protection mechanism of *C. ohadii* is different from that of the model alga *C. reinhardtii*. Differences between the PI protection mechanisms and structural properties of the two algae grown under LL and HL conditions are listed. (+) present, (-) absent, (?) not known. There is a large fluorescence quenching in HL *C. ohadii* but it is different from the classical NPQ described for *C. reinhardtii*.

|  | *C. ohadii* | *C. reinhardtii* |
| --- | --- | --- |
| Grow at 2500 $\mu\text{E m}^{-2}\text{s}^{-1}$ | + | - |
| PSII |  |  |
| PSI |  |  |
| Chl *a/b* | 5 (LL) >10 (HL) | ~3-4 |
| Xanthophyll cycle | + | + |
| PSII repair cycle | + | + |
| LHCSR | - | + |
| PSBS | - | - (transient) |
| CBR, ELIP | ++++ | +++ |
| PTOX | +/- | + |
| PSII protein phosph. | - | + |
| PSI-CEF in HL | ? | + |
| State transitions | - | + (up to 80%) |
| NPQ | ++ non-classic | ++ |

First, *C. ohadii* and its close relative *C. sorokiniana* were confirmed to be better protected against PI compared to other photosynthetic organisms. The two species thrived at a HL intensity of 2500 µE m^-2^s^-1^, where several other green algae and a model cyanobacteria were bleached. As the results of their extreme HL habitats, *C. ohadii* and *C. sorokiniana* developed most efficient PI protection mechanism. The differences between the PI protection mechanism of *C. ohadii* and the model alga *C. reinhardtii* are listed in Table I and discussed below.

### Lack or very low expression of LHCSR and psbS in HL cells

LHCSR and psbS are two key stress-related proteins in the PI protection processes of *C. reinhardtii* and plants. These proteins sense the acidification of the lumen as the result of robust PSII activity at HL and activate the NPQ and the xanthophyll cycle protection mechanisms ^46,47,49^. LHCSR has been shown to be required for the effective PI protection of *C. reinhardtii* and plants ^2^. Since *C. ohadii* and *C. sorokiniana* are more resistant to PI than *C. reinhardtii*, it was surprising to find no LHCSR homolog in their genomes. Therefore, the efficient PI protection must be performed without the activity of LHCSR members. Moreover, these *Chlorella* strains eliminated the LHCSR genes throughout their evolution while developing more effective PI protection processes. In contrast, two psbS genes were present in their genomes and in the case of *C. ohadii*, the corresponding transcripts were previously detected ^23^. However, psbS was not detected by MS and to much less amount than in *C. reinhardtii* by immunodetection (Fig 4). Therefore, psbS is either entirely absent or present in minor undetectable amount in HL thylakoids. Accordingly, its function in sensing lumen acidification and initiating PI protection processes is limited.

A seemingly appropriate PI protection initiator candidate that emerged in the current study was CBR, which bears a structure similar to that of LHCSR and psbS (Fig 4) and significantly accumulated in the LHC, where both LHCSR and psbS are located in *C. reinhardtii* and plants ^2^. A search for conserved glutamates facing the lumen that may sense acidification, revealed one in CBR, compared to two in psbS (Fig 4d and S7). A more detailed analysis of CBR will still be necessary to determine if it senses pH changes in the lumen and initiates PI protection.

LHCSR and psbS evolved in photosynthetic organisms as important players in the developing of an efficient PI protection. It remains to be determined when *C. ohadii* and *C. sorokiniana* lost the LHCSR genes and why there is no need for LHCSR and psbS in the most efficient PI-protecting mechanism known thus far in nature.

### Advantage of the elimination of PSII antenna under HL

The major structural difference between LL and HL cells is the change in the PSII at HL (Table I). A large PSII-LHCII complex, similar to that of *C. reinhardtii*, is present in LL cells, while a core PSII lacking both the inner and outer antenna is prominent in HL cells. While *C. reinhardtii* also showed fluctuations in the antenna size under HL, the extreme reduction of both outer and inner antenna, as shown in this work, was unique to *C. ohadii* and shown to be important for PI protection. The drastic reduction of the antenna size significantly reduces the amount of light that is funneled to the reaction center and reduces the amount of harmful ROS and accordingly, the level of PI. In addition, the PSII repair cycle may work more rapidly with the naked complex than with the full-size LL structure. On the other hand, LHCII is a major site of scavenging and quenching of ROS, the harmful side products of overexcitation under HL ^2^. It is possible that the large amount of xanthophylls that accumulates in the remaining LHCII molecules and the lipid moiety, scavenges and sequesters the radicals (Fig. 7 and Table S3). In this way, the enormous amount of lutein and zeaxanthin replaces the scavenging function LHCII plays in other green algae and plants. Alternatively, previous works have suggested a role for CBR in protection against PI ^33,41,43^. The large amount of CBR that accumulates under HL conditions strongly suggests a scavenging function in PI protection. Taken together, in order live in the desert biological sand crust, *C. ohadii* developed PI protection mechanisms, including elimination of the PSII antenna. While serving as an advantage for the rapid PSII repair process, the building of the large complex and degrading the LHCII proteins is time- and energy-consuming and is likely disadvantageous if performed on a daily schedule. Indeed, the present work showed that the adaptation of the PSII structure is a prolonged process that probably occurs on a seasonal basis (Fig. 7).

### Lack of state transitions and PSII protein phosphorylation

The process of state transitions guarantees correct balancing of the absorbed light between the two photosystems in order to optimize the photosynthetic electron flow ^12^. In addition, it has been proposed to be important to PI protection through its reduction of the PSII antenna when PSII is overexcited and over-reduces the plastoquinone pool. This process has been shown to be a key contributor to PI protection in *C. reinhardtii*, by reducing the LHCII light absorption under HL by up to 80%^30^. As *C. ohadii* is more resistant to HL than *C. reinhardtii*, it was somewhat surprising to find out that state transition is very limited under LL and does not occur at all in HL cells. While PSII in HL cells is stripped of LHCII, and therefore, no further detachment and transition of LHCII to PSI can be expected, it is less clear why very limited state transitions occurred in LL cells, where the PSII is surrounded by LHCII (Fig 2). The lack of state transitions was strengthened by the absence of phosphorylated LHCII and PSII proteins. Interestingly, dephosphorylation of PSII and LHCII proteins during PI of *C. reinhardtii* and the loss of state transitions while retaining activity of the cyclic electron pathway have been reported long time ago ^50,51^. The presence of stt7 and stl1 homologs in the genome, suggests that state transitions was eliminated during *C. ohadii* evolution and perhaps during the development of its efficient PI protection mechanisms. The lack of state transitions, phosphorylated PSII-LHCII proteins, LHCSR and psbS and the formation of the core-PSII complex at HL, are all unique features of *C. ohadii* that significantly differentiate its PI protection mechanism from that described for *C. reinhardtii*. It will be interesting to determine how the exclusion of these functions benefited the development of the more efficient PI protection mechanism.

### The change of PSII structure in the acclimation to HL is likely a seasonal and not diurnal process

The summer day in the Israeli desert biological sand crust begins at sunrise, with 3-4 h of relatively LL intensity and humid air as the result atmospheric moisture (dew and fog), which are excellent conditions for photosynthesis^52^. Then, there are 8-12 h of extreme HL and dryness, i.e., the period of PI, followed by the night hours. While it is possible that some of the PI protection processes, such as fluorescence quenching and activation of the xanthophyll cycle, occur on a diurnal cycle, the reduction of the PSII antenna at HL and its rebuilding at LL seems to be too energy-consuming to be advantageous if performed on a daily basis. Indeed, when following the adaptation of LL cells to HL, xanthophylls, lutein and zeaxanthin accumulation began shortly after the onset of HL, while the reduction in LHCII-PSII, the accumulation of core PSII and the accompanying decrease of chlorophyll *b*, begin only after 8 h of HL (Fig 7). This suggests a delay in the signal transduction and activation of the transformation of LHCII-PSII to the core PSII form which characterizes HL cells. This lag may have developed to prevent diurnal fluctuation of the PSII structure, and to ensure a more seasonal change. In summary, the HL structure of the core PSII is likely established during the initial days of the summer, when there are more hours of HL intensities, and is maintained throughout the long summer. Transformation to the LL PSII-LHCII likely occurs when autumn arrives, when the day become shorter and light intensity is reduced. Since the cells rapidly multiply, it will be interesting to determine whether the LHCII proteins and chlorophylls are degraded during adaptation from LL to HL, or if they are diluted in the multiplying cells. Further studies focusing on signal transduction and the control of gene expression during cell adaptation to LL and HL conditions will be necessary to shed light on these topics.

### Conclusions

This work showed that *C. ohadii*, a green alga that thrives under HL intensities which trigger PI and eventual death in other algae, developed a unique PI protection mechanism that in several features significantly differed from the well-established mechanism of the model green alga *C. reinhardtii*. More specifically, *C. ohadii* showed no state transitions, PSII and LHCII phosphorylation or expression of stress proteins LHCSR and psbS. In their place, the alga featured core, LHC-less PSII, overexpression of CBR and accumulation of high concentrations of protective xanthophylls in the remaining LHCII and the lipid moiety. Together, these unique features provided for PI protection that outperformed the efficacy of the mechanism of the model green alga *C. reinhardtii* and other photosynthetic organisms.

## Methods

### Cells and growth conditions

*C. ohadii* was grown in Tris-acetate-phosphate (TAP) medium at 28 °C, as previously described ^25^. Cultures were grown under white light of either 2000 (HL) or 100 μE m^-2^s^-1^ (LL). To achieve full acclimation to HL, the cells were grown in HL for 6 generations and maintained below OD_750nm_=0.7 to prevent self-shading. The other strains were grown in TAP medium, under LL conditions. *D. salina* were grown in BG11 medium supplemented with 1.5 M NaCl. *S. elongatus* was grown in BG11 medium.

### State transitions

States I and II were induced by incubating cells 20 min at 100 µE m^-2^s^-1^ with 50 µM DCMU or 20 µM FCCP, respectively^53^. The 77K fluorescence spectra was done as described^25^.

### Chlorophyll and protein concentrations

Chlorophyll and protein concentrations were measured as described ^25^.

### Large- and low-scale thylakoid preparations

Thylakoids were extracted as previously described ^25^.

### Protein immunoblotting

Proteins were immunoblotted as previously described ^54^. The following antibodies, purchased from Agrisera, were used: lhcb1 - AS01 004, lhcb2 - AS01 003, lhcb4 - AS04 045, lhcb5 - AS01 009, lhcb6 - AS01 010, lhca2 - AS01 006, psbA - AS11 1786, psbC - AS11 1787, psbD - AS06 146, psaA - AS06 172, petC - AS08 330, psbS - AS09 533. Antibodies recognizing threonine-phosphorylated LHCII proteins were purchased from Cell Signalling Technology – 9381S. The identification of the *C. ohadii* LHCII proteins that reacted with LHC antibodies raised against the LHC subunits of *A. thaliana* was as described before ^25^.

### Native PAGE

Thylakoid membranes were solubilized for 20 min, at 25 °C, in 2% β-dodecyl maltoside (βDDM), 0.7% n-octyl-β-d-glucoside (OG) and 1% sodium dodecyl sulfate (SDS), and then cleared by centrifugation (12,000 *g*, 10 min). Thereafter, the extracted protein complexes were fractionated on by gradient 3-12% acrylamide-PAGE and photographed. The fluorescence of the fractionated complexes was measured with a *Typhoon FLA 7000*, at an excitation wavelength of 532 nm.

### Sucrose density gradient separation of chlorophyll protein complexes

Thylakoid membranes were solubilized for 10 min, at 4 °C, in 1% -DDM and then cleared by centrifugation (12,000 *g*, 10 min). Thereafter, the extracted protein complexes were fractionated over a 4-45% sucrose density gradient in 25 mM MES (pH 6.5) buffer, by centrifugation at 100,000 *g*, for 20 h, using a SW-41 rotor.

### Mass spectrometry

Label-free thylakoid membranes and isolated photosynthetic complexes of LL and HL C. *ohadii* were prepared as described ^25^. Liquid chromatography, tandem mass spectrometry (LC-MS/MS) and data analysis were also performed as described ^25^, with the following modifications: Raw data were processed using MaxQuant software (1.6.3.4)^55^ to obtain intensity-based absolute quantification (iBAQ) values. iBAQ values correspond to the abundance of a given protein in the sample. Therefore, the iBAQ value for a given protein is determined by calculation of the sum of all identified peptide intensities divided by the number of theoretically observable peptides ^56^. Relative iBAQ (riBAQ), which is the abundance of the protein relative to the total protein amount, was calculated for a given protein by dividing its iBAQ value by the sum of the iBAQ values of all proteins in the sample. riBAQ values are between 0 and 1 and are equivalent to normalized molar intensity ^35,36^. This means that if a given protein has a riBAQ value of 0.2, its protein molarity comprises 20% of the total protein molarity in the sample. To calculate the relative abundance in percent (%), riBAQ values were multiplied by 100.

### Carotenoid analysis

Frozen thylakoid and photosynthetic complexes obtained by sucrose density gradient fractionation were extracted in 8 ml hexane:acetone:ethanol (50:25:25, v/v), and then saponificated for 5 min in 80% KOH and two extractions with hexane. Carotenoid analysis by HPLC using a reverse phase C18 column was carried out as previously described^57^. Further manipulations and data analysis were performed as described ^25^.

### ROS analysis

Dark-adapted thylakoids and photosynthetic complexes were spotted in serial dilutions on PVDF membranes under dim light, using a dot-plot apparatus. For intact cells, the ROS detection reactions were conducted in Eppendorf microtubes and then spotted manually onto Whatman 3 filter paper. A solution containing 5 mM of nitro blue tetrazolium (NBT) or 3,3′-diaminobenzidine (DAB) in a buffer containing HEPES 15 mM pH 8.0, NaCl 15 mM, MgCl_2_ 5 mM and sucrose 0.3 M, was added and the membrane was then illuminated with a light intensity of 1000 µE m^-2^s^-1^, for 10 min. The membranes were then washed and chlorophyll was extracted using 100% methanol. Quantitation was performed using the Image J software and the linear range of the color density was used to determine ROS levels.

### Protein structural modeling

Homology modeling of CBR and psbS structures was performed using Swiss-Model^58^. The suggested structure was also verified with the *C. Sorokiniana* CBR obtained from the alphafold protein structure database^59,60^

## Funding

Funding was provided by a “Nevet” grant from the Grand Technion Energy Program (GTEP) and a Technion VPR Berman Grant for Energy Research.

We thank Aaron Kaplan for providing *C. ohadii* and *C. sorokiniana* cells, and Liat Young, Iftach Yacoby and Nir Keren for help in experiments and valuable advices and discussions.

## Author contribution

GL and GS designed the research. GL, MY and GS performed the research. AM, MC, YT and JH performed the carotenoids analysis. GL, GS and NA wrote the article.

## Conflict of interests

The authors declare no competing financial interests.

## Data availability statement

All original data and material information requests can be made to either Gadi Schuster at or Guy Levin at.

## Supporting information

Additional supporting information can be found in the online version of this article (Tables S1-3 and Figures S1-8).

## Notes

### Competing Interest Statement

The authors have declared no competing interest.

