## Supplemental Tables and Figures for "A desert green alga that thrives at extreme high-light intensities using a unique photoin-hibition protection mechanism"

| | | Relative of Total protein in fraction (%) | | | | Relative to D2 (%) | | | |
|---|---|---|---|---|---|---|---|---|---|
| **Uniprot I.D** | **Protein** | **HL LHCII** | **HL PSII Core** | **LL LHCII** | **LL PSII-LHCII** | **HL LHCII** | **HL PSII Core** | **LL LHCII** | **LL PSII-LHCII** |
| W8SYD4 | psbD (D2) | 0.2 | **4.1** | 0.2 | **3.8** | 100 | **100** | 100 | **100** |
| W8SIR2 | psbA (D1) | 0.2 | **2.8** | 0.04 | **1.2** | 106 | **67** | 20 | **31** |
| W8TIK4 | psbB (CP 47) | 1.0 | **8.3** | 0.4 | **4.6** | 531 | **201** | 236 | **122** |
| W8SUG2 | psbC (CP43) | 0.3 | **6.8** | 0.5 | **3.7** | 133 | **165** | 279 | **98** |
| W8SIT0 | psbH | 0.3 | **6.0** | 0.7 | **6.0** | 165 | **146** | 352 | **160** |
| W8SU91 | psbE | 0.2 | **2.6** | 0.1 | **1.3** | 112 | **62** | 60 | **36** |
| W8SIQ3 | psbF | 0.5 | **0.6** | 0.4 | 0.2 | 250 | **15** | **212** | 6 |
| A0A2P6TWR3 | psbO | **0.4** | 0.3 | 0.2 | **0.5** | **196** | 8 | 118 | **13** |
| A0A2P6U3U0 | psbP | **0.5** | 0.4 | 0.3 | **1.4** | **259** | 10 | **185** | **37** |
| A0A2P6TGD3 | psbQ | 0.04 | **0.1** | 0.004 | **0.4** | 19 | **1.8** | **19** | **9.7** |
| A0A2P6TSU7 | psbR | 0.0002 | **0.02** | 0.001 | **0.1** | 0.2 | **0.5** | 0.7 | **3.4** |
| A0A2P6TFK0 | psb27 | ND | **0.2** | ND | 0.001 | ND | **4.8** | ND | 0.03 |
| A0A2P6TPR3 | psb28 | 0.0 | **0.1** | 0.0 | 0.0 | 10.5 | **3.3** | 6.9 | ND |
| A0A2P6TTG5 | CP 26 | **1.3** | 0.5 | 2.4 | 4 | **672** | 11 | 1292 | 107 |
| A0A2P6U4D0 | CP 29 | **0.6** | 0.2 | 0.7 | 2.2 | **295** | 4 | 357 | 58 |
| A0A2P6TDA6 | LHCBM2 | **6.4** | 0.4 | 10.1 | 4.4 | **3254** | 11 | 5428 | 117 |
| A0A2P6U146 | LHCBM4 | **3.5** | 0.2 | 15.4 | 3.8 | **1778** | 5 | 825 | 100 |
| W8SUC6 | psbN | 1.0 | 0.2 | **0.2** | ND | **515** | 4.2 | **97** | ND |
| A0A2P6TKP3 | psbP (chloroplastic) | 0.009 | **0.02** | **0.005** | 0.0003 | 4.5 | **0.5** | **2.7** | 0.01 |
| A0A2P6TL23 | psbP containing | **0.002** | 0.0005 | ND | ND | **1** | 0.01 | ND | ND |
| A0A2P6TXN8 | psbP containing | **0.02** | 0.006 | **0.005** | ND | **8.6** | 0.1 | **2.8** | ND |
| A0A2P6U5H8 | HCF 136 | **0.2** | 0.2 | **0.03** | 0.002 | **120** | 4.7 | **17.2** | 0.06 |
| A0A2P6U2Y6 | LPA | **0.004** | ND | **0.0008** | ND | **1.8** | ND | **0.4** | ND |
| A0A2P6TZ87 | LPA (2) | **0.009** | 0.003 | **0.004** | 0.002 | **4.5** | 0.07 | **1.9** | **0.06** |
| A0A2P6TD59 | LPA (3) | **0.06** | 0.004 | **0.03** | 0.0006 | **28** | 0.1 | **15** | 0.01 |
| A0A2P6TPG7 | LPA (4) | ND | **0.0005** | **0.0008** | ND | ND | **0.01** | **0.45** | ND |

**Table S1. Mass spectrometry detection and quantification of PSII and LHCII related proteins in photosynthetic complexes of LL and HL *C. ohadii*.** The values to the left half of the table correspond to the relative abundance of each protein in percent (%) of total protein molarity in each sample. i.e., if the total protein concentration equals xM (=100%), and the relative abundance value of a given protein is 1%, than the concentration of this protein is 0.01xM (see materials and methods). Numbers to the right half of the table are the abundance of a protein as compared to the abundance of D2, in percent (%). It was calculated by dividing a given protein riBAQ value by the riBAQ value of D2 and multiplying it by 100.

| Uniprot I.D | Protein | HL LHCII | HL PSII Core | Thylakoids | LL LHCII | LL PSII-LHCII | Thylakoids |
|---|---|---|---|---|---|---|---|
| **A0A2P6TNM2** | **CBR** | **1.8** | **0.2** | **0.1** | **nd** | **nd** | **nd** |
| A0A2P6U2C2 | ELIP1 | 0.04 | 0.003 | 0.02 | nd | nd | 0.00003 |
| A0A2P6TQZ2 | ELIP2 | 0.02 | 0.0007 | 0.01 | 0.0006 | nd | 0.001 |
| A0A2P6TDF0 | LM ELIP | 0.24 | 0.08 | 0.1 | 0.8 | 0.002 | 0.05 |
| A0A2P6U416 | HLIP (OHP1) | 0.16 | 0.03 | 0.05 | 0.03 | 0.001 | 0.03 |
| A0A2P6TTD5 | HLIP (OHP2) | 0.14 | 0.02 | 0.005 | 0.03 | nd | 0.004 |
| A0A2P6TJB9 | psbS | nd | nd | nd | nd | nd | nd |
| W8SYD4 | psbD (D2) | 0.2 | 4.1 | 1.2 | 0.2 | 3.8 | 2.2 |

**Table S2. Mass spectrometry detection and quantification of LHC-like proteins in photosynthetic complexes and thylakoids of LL and HL *C. ohadii* cells.** The values correspond to the relative abundance of each protein in percent (%) of total protein molarity in each sample. i.e. if the total protein concentration equals xM (=100%), and the relative abundance value of a given protein is 1%, than the concentration of this protein is 0.01xM (see materials and methods.) D2 value is given as reference.

| Time (h) | Neoxanthin [µg/cell (x10-8)] | Violaxanthin [µg/cell (x10-8)] | Antheraxanthin [µg/cell (x10-8)] | Zeaxanthin [µg/cell (x10-8)] | β-Carotene [µg/cell (x10^-8)] | Lutein [µg/cell (x10-8)] | α-Carotene [µg/cell (x10-8)] | Total [µg/cell (x10-8)] | Car/Chl (µg/µg) |
|---|---|---|---|---|---|---|---|---|---|
| 0 | 0.35 | 0.43 | 0.13 | 0.04 | 1.00 | 2.40 | 0.08 | 4.44 | 0.12 |
| 1 | 0.52 | 0.67 | 0.42 | 0.53 | 1.69 | 4.65 | 0.07 | 8.55 | 0.23 |
| 3 | 0.79 | 0.46 | 0.65 | 3.23 | 2.24 | 6.70 | 0.07 | 14.14 | 0.37 |
| 5 | 0.69 | 0.34 | 1.14 | 3.39 | 2.11 | 7.33 | 0.06 | 15.06 | 0.51 |
| 8 | 0.18 | 0.37 | 1.02 | 2.01 | 1.54 | 5.06 | 0.04 | 10.22 | 0.47 |
| 16 | 0.18 | 0.38 | 0.93 | 1.76 | 1.69 | 4.81 | 0.06 | 9.81 | 0.52 |
| 24 | 0.15 | 0.65 | 1.26 | 1.89 | 2.39 | 6.12 | 0.06 | 12.53 | 0.66 |

| Complex | Cells | Neoxanthin (carotenoids/cell – arbitrary) | Violaxanthin (carotenoids/cell – arbitrary) | Antheraxanthin (carotenoids/cell – arbitrary) | Zeaxanthin (carotenoids/cell – arbitrary) | β-Carotene (carotenoids/cell – arbitrary) | Lutein (carotenoids/cell – arbitrary) | α-Carotene (carotenoids/cell – arbitrary) | Total (carotenoids/cell – arbitrary) | Car/Chl (µg/µg) |
|---|---|---|---|---|---|---|---|---|---|---|
| LHCII | LL | 15.5 | 14.0 | 2.0 | 3.6 | 0.2 | 86.6 | 0.0 | 121.8 | 0.1 |
| | HL | 12.5 | 38.8 | 58.3 | 100.2 | 10.6 | 247.4 | 0.2 | 469.9 | 0.5 |
| PSII-LHCII | LL | 14.4 | 11.1 | 2.5 | 1.4 | 34.1 | 66.9 | 1.9 | 132.8 | 0.1 |
| PSII core | HL | 0.4 | 0.3 | 0.7 | 1.0 | 8.0 | 2.8 | 0.3 | 13.5 | 0.03 |
| Free Pigment | LL | 34.3 | 151.5 | 17.8 | 33.8 | 15.6 | 427.0 | 1.3 | 685.8 | 0.7 |
| | HL | 28.2 | 268.8 | 424.7 | 618.9 | 73.1 | 1717.8 | 0.0 | 3147.3 | 3.1 |
| PSI-LHCI | LL | 23.4 | 5.8 | 0.0 | 0.0 | 22.2 | 91.9 | 5.7 | 149.0 | 0.15 |
| | HL | 16.0 | 3.8 | 19.4 | 13.1 | 39.8 | 93.6 | 2.2 | 187.9 | 0.2 |

**Table S3. Carotenoids accumulate in HL cells. (a)** LL cells were transferred to HL and samples were taken at the indicated times following the transfer to HL and their carotenoids analyzed. **(b)** Accumulation of carotenoids in photosynthetic complexes and the free pigment fraction.

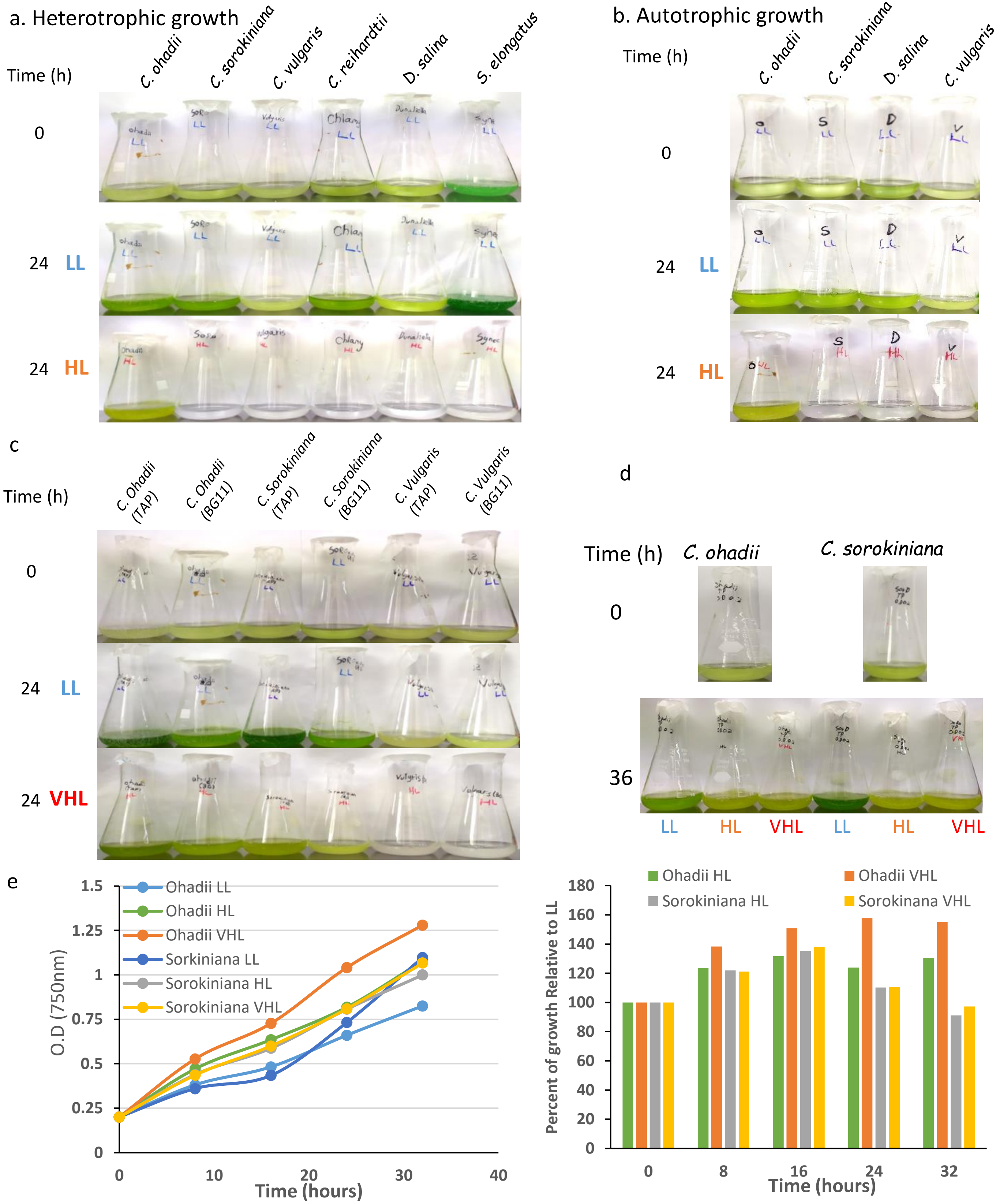


**Figure S1. *C. Ohadii* thrives where other organism do not survive. (a)** Experiment performed in carbon containing growth medium. *S. elongatus* in BG11 medium, *D. salina* in BG11+ 1.5M $NaCl_2$, all other organisms in TAP medium. All cultured started at $O.D_{750}$ =0.14. **(b)** same as (a), but without carbon source in growth mediums. **(c)** Same as (a), but each organism tested in 2 different growth mediums (TAP and BG11). All cultures started at $O.D_{750}$ = 0.2. **(d)** Similar to (a), but with no carbon source (TP medium) and starting $O.D_{750}$ = 0.2. **(e)** growth curves (left) and comparison of growth compared to LL cells (right) of *C. ohadii* and *C. sorokiniana* under LL, HL or VHL conditions. LL, HL and VHL are 100, 2000 and 2500 µmols photons $m^{-2}s^{-1}$. Note that the results for *C. sorokiniana* were not conclusive. In some experiments it was photoinhibited, as shown in panels a and b, while in others its growth was not affected, as shown in panels c and d.

a

**PTOX**

| | Description | Scientific Name | Max Score | Total Score | Query Cover | E value | Per. Ident | Acc. Len | Accession |
|---|---|---|---|---|---|---|---|---|---|
| ☐ | plastid terminal oxidase [Chlorella sorokiniana] | Chlorella sorokiniana | 389 | 389 | 90% | 9e-133 | 54.69% | 482 | PRW56883.1 |

**stt7**

| | Description | Scientific Name | Max Score | Total Score | Query Cover | E value | Per. Ident | Acc. Len | Accession |
|---|---|---|---|---|---|---|---|---|---|
| ☑ | Serine threonine- kinase chloroplastic [Chlorella sorokiniana] | Chlorella sorokiniana | 460 | 460 | 72% | 8e-153 | 47.89% | 694 | PRW20521.1 |
| ☑ | serine threonine- kinase chloroplastic-like [Chlorella sorokiniana] | Chlorella sorokiniana | 199 | 199 | 44% | 1e-53 | 36.02% | 1037 | PRW58226.1 |
| ☑ | Serine threonine- kinase chloroplastic [Chlorella sorokiniana] | Chlorella sorokiniana | 190 | 190 | 51% | 6e-53 | 33.72% | 506 | PRW05929.1 |
| ☑ | cell division control 2-like protein [Chlorella sorokiniana] | Chlorella sorokiniana | 63.2 | 63.2 | 30% | 1e-10 | 29.48% | 424 | PRW20833.1 |
| ☑ | serine threonine kinase [Chlorella sorokiniana] | Chlorella sorokiniana | 61.6 | 61.6 | 33% | 3e-10 | 26.47% | 363 | PRW57463.1 |

**stl1**

| | Description | Scientific Name | Max Score | Total Score | Query Cover | E value | Per. Ident | Acc. Len | Accession |
|---|---|---|---|---|---|---|---|---|---|
| ☑ | Serine threonine- kinase chloroplastic [Chlorella sorokiniana] | Chlorella sorokiniana | 467 | 467 | 82% | 1e-161 | 57.45% | 506 | PRW05929.1 |
| ☑ | Serine threonine- kinase chloroplastic [Chlorella sorokiniana] | Chlorella sorokiniana | 191 | 191 | 82% | 1e-53 | 34.64% | 694 | PRW20521.1 |
| ☑ | serine threonine- kinase chloroplastic-like [Chlorella sorokiniana] | Chlorella sorokiniana | 164 | 164 | 70% | 2e-43 | 32.61% | 1037 | PRW58226.1 |
| ☑ | Serine threonine- kinase 36 [Chlorella sorokiniana] | Chlorella sorokiniana | 60.1 | 60.1 | 12% | 1e-09 | 45.45% | 1728 | PRW61305.1 |
| ☑ | MAP3K epsilon kinase 1-like isoform X2 [Chlorella sorokiniana] | Chlorella sorokiniana | 57.8 | 57.8 | 43% | 6e-09 | 23.61% | 1448 | PRW61538.1 |

b

| Uniprot I.D | Protein | HL LHCII | HL PSII Core | Thylakoids | LL LHCII | LL PSII-LHCII | Thylakoids |
|---|---|---|---|---|---|---|---|
| A0A2P6TS42 | PTOX | 0.02 | 0.0008 | 0.0014 | 0.01 | nd | 0.0003 |
| A0A2P6TD00 | stt7 | 0.002 | 0.0006 | 0.002 | 0.002 | 0.00004 | 0.002 |
| A0A2P6TBJ9 | stl1 | nd | nd | 0.003 | nd | 0.0003 | 0.005 |

**Figure S2. PTOX, stt7 and stl1 kinases homologs are expressed in *C. ohadii*. (a)** NCBI protein BLAST analysis of PTOX2 (uniport I.D - A8IEF7), stt7 (Q84V18) and stl1 (Q84V17) of *C. reinhardtii* vs. the proteome of *C. Sorokiniana*. Hits highlighted by a black box were implicated as the homologs of the corresponding protein in *C. ohadii*. **(b)** Mass spectrometry relative intensity based absolute quantification (riBAQ) analysis of the relative abundances of the selected homologs in thylakoid membranes and photosynthetic complexes of *C. ohadii*. The values correspond to the relative abundance of each protein in percent (%) of total protein molarity in each sample. i.e. if the total protein concentration equals xM (=100%), and the relative abundance value of a given protein is 1%, than the concentration of this protein is 0.01xM (see materials and methods).

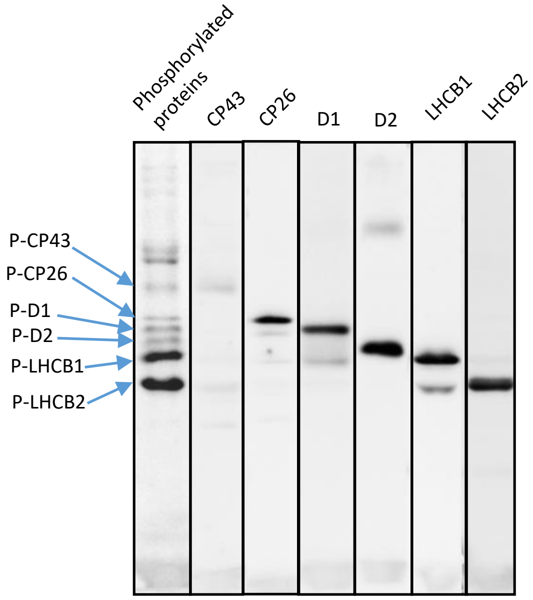


**Figure S3. Identification of the *C. reinhardtii* phosphorylated thylakoid proteins*.*** Phospho-threonine/tyrosine antibody was used to detected phosphorylated thylakoid proteins in *C. reinhardtii* cells locked in state II (1$^{st}$ lane to the left). Immuno-blotting of photosynthetic proteins with specific antibodies disclosed the migration of each on SDS-PAGE as compared to the phospho-proteins (lanes 2-6 from the left).

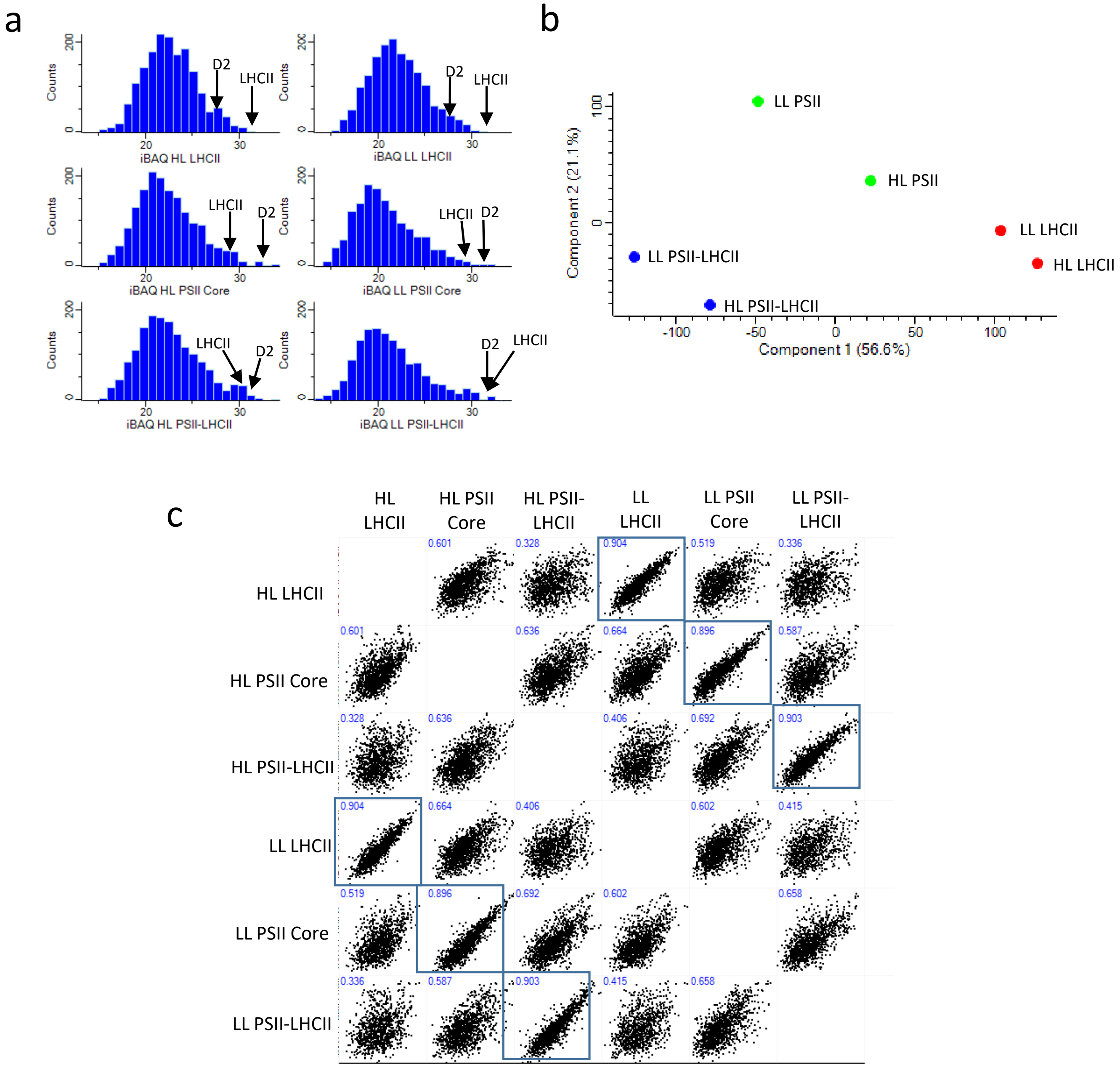


d

| | Total proteins: 2291 | | | Total proteins: 2163 | | |
|---|---|---|---|---|---|---|
| | HL LHCII | HL PSII Core | HL PSII-LHCII | LL LHCII | LL PSII Core | LL PSLHCII |
| Total proteins in sample: | 1718 | 1704 | 1652 | 1664 | 1448 | 1411 |
| Number of max abundance proteins: | 1221 | 301 | 769 | 1200 | 326 | 637 |
| Dynamic Range: | 6.4% - 0.00004% | 8.3% - 0.00001% | 8.2% - 0.00002% | 15.4% - 0.00002% | 8.5% - 0.00003% | 6% - 0.00001% |

**Figure S4. Mass spectrometry analysis data quality check. (a)** Histogram of protein abundance in each sample of the photosynthetic complexes of LL and HL cells. Values in X-axis correspond to the log transformed [log2(x)] iBAQ value of any given protein. Values on Y axis correspond to the number of proteins detected with that iBAQ value. **(b)** Principal component analysis of the analyzed samples. **(c)** Pearson correlation between samples. **(d)** Number of proteins detected in each sample (top), the number of proteins which are mostly abundant in the given sample (middle) and the dynamic range in each sample (bottom). i.e – dynamic range is the range between the riBAQ values (see materials and methods) of the most abundant and most scarce proteins in the sample.

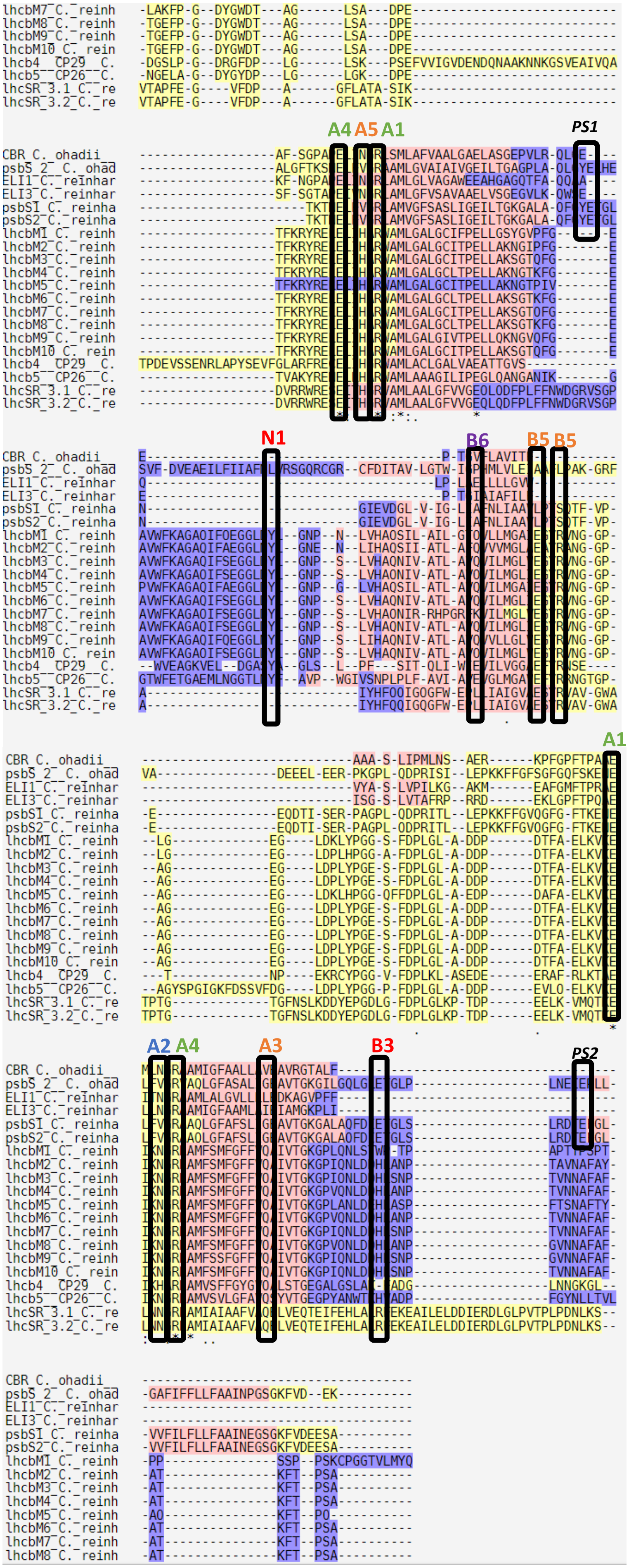


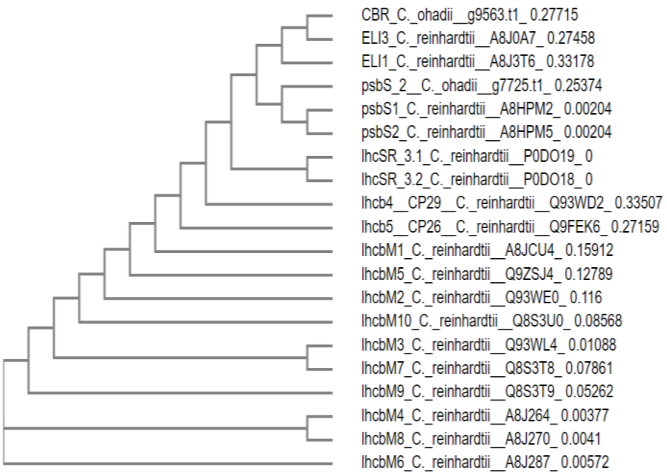


**Figure S5. CBR is able to bind an extra chlorophyll compared to psbS. (a)** Transmembrane protein multiple sequence alignment of *C. ohadii* CBR and psbS(2) homologs vs psbS1, psbS2, ELIP3, ELIP1, lhcbM 1-10, CP26, CP29, lhcSR 3.1 and lhcSR 3.2 of *C. reinhardtii*. Conserved residues which bind chlorophyll *a* (An) and *b* (Bn), and neoxanthin (N1) are indicated. Note that psbS is capable of binding two chlorophyll *a* molecules (A1 and A4) while CBR is capable of binding an additional chlorophyll (A1, A2 and A4). The color of each pigment name (A1, A2 etc) is directly related to the proteins that contain the residues for its binding. Green – residue conserved in all tested proteins. Blue – conserved in all except psbS. Orange – conserved in all but psbS, CBR and ELIP1/3. Red – conserved only in CP26/29 and lhcbm 1-10. Purple – conserved only in lhcbM 1-10. ps – proton sensing residues. **(b)** sequence based phylogenetic tree. Note that although CBR contains only three transmembrane alpha helices, similar to lhcSR and lhcbM, it is more closely related to the four helices containing psbS.

a

(CBR) *C. sorokiniana* to *C. ohadii*

| | Description | Scientific Name | Max Score | Total Score | Query Cover | E value | Per. Ident | Acc. Len | Accession |
|---|---|---|---|---|---|---|---|---|---|
| ☑ | g9563.t1 (CBR) | | 291 | 291 | 100% | 6e-102 | 84.10% | 193 | Query_47799 |
| ☑ | g3146.t1 | | 101 | 101 | 53% | 2e-28 | 50.96% | 104 | Query_41148 |
| ☑ | g1315.t1 | | 66.2 | 66.2 | 55% | 8e-14 | 41.86% | 230 | Query_39243 |
| ☑ | g10714.t1 | | 47.8 | 47.8 | 46% | 5e-08 | 37.11% | 96 | Query_48998 |
| ☑ | g7554.t1 | | 38.5 | 38.5 | 47% | 4e-04 | 35.09% | 191 | Query_45707 |

b

(CBR) *C. ohadii* to *C. reinhardtii*

| | Description | Scientific Name | Max Score | Total Score | Query Cover | E value | Per. Ident | Acc. Len | Accession |
|---|---|---|---|---|---|---|---|---|---|
| ☑ | uncharacterized protein CHLRE_09g394325v5 [Chlamydomonas reinhardtii] | Chlamydomonas reinhardtii | 138 | 138 | 89% | 3e-41 | 45.51% | 197 | XP_0016946[illegible]1.1 |
| ☑ | uncharacterized protein CHLRE_09g393173v5 [Chlamydomonas reinhardtii] | Chlamydomonas reinhardtii | 74.3 | 74.3 | 54% | 8e-17 | 34.68% | 175 | XP_001694751.1 |
| ☑ | uncharacterized protein CHLRE_07g320400v5 [Chlamydomonas reinhardtii] | Chlamydomonas reinhardtii | 70.5 | 70.5 | 51% | 3e-15 | 40.52% | 190 | XP_042922535.1 |
| ☑ | uncharacterized protein CHLRE_08g384650v5 [Chlamydomonas reinhardtii] | Chlamydomonas reinhardtii | 68.9 | 68.9 | 56% | 2e-14 | 40.50% | 230 | XP_001694210.2 |
| ☑ | uncharacterized protein CHLRE_16g679250v5 [Chlamydomonas reinhardtii] | Chlamydomonas reinhardtii | 64.7 | 64.7 | 54% | 3e-13 | 43.12% | 176 | XP_001695978.1 |

CBR homolog in C. reinhardtii: **ELIP3** (uniport I.D - A8J0A7)

c

(psbS) *C. sorokiniana* to *C. ohadii*

| | Description | Scientific Name | Max Score | Total Score | Query Cover | E value | Per. Ident | Acc. Len | Accession |
|---|---|---|---|---|---|---|---|---|---|
| ☑ | g7724.t1 (PSBS 1) | | 427 | 568 | 55% | 2e-135 | 86.31% | 1814 | Query_22587 |
| ☑ | g7726.t1 (PSBS 3) | | 388 | 388 | 41% | 1e-134 | 92.86% | 259 | Query_22589 |
| ☑ | g7725.t1 (PSBS 2) | | 373 | 432 | 55% | 2e-128 | 74.75% | 300 | Query_22588 |
| ☑ | g7554.t1 | | 82.4 | 129 | 38% | 1e-18 | 35.62% | 191 | Query_22405 |
| ☑ | g7763.t1 | | 56.2 | 56.2 | 38% | 1e-08 | 23.93% | 620 | Query_22629 |

(psbS[1]) *C. ohadii* to *C. reinhardtii*

| | Description | Scientific Name | Max Score | Total Score | Query Cover | E value | Per. Ident | Acc. Len | Accession |
|---|---|---|---|---|---|---|---|---|---|
| ☑ | uncharacterized protein CHLRE_01g016750v5 [Chlamydomonas reinhardtii] | Chlamydomonas reinhardtii | 206 | 206 | 10% | 6e-60 | 62.30% | 245 | XP_001689923.1 |
| ☑ | uncharacterized protein CHLRE_01g016600v5 [Chlamydomonas reinhardtii] | Chlamydomonas reinhardtii | 206 | 206 | 10% | 6e-60 | 62.30% | 245 | XP_001689476.1 |
| ☑ | uncharacterized protein CHLRE_03g146147v5 [Chlamydomonas reinhardtii] | Chlamydomonas reinhardtii | 80.5 | 80.5 | 8% | 8e-16 | 37.75% | 307 | XP_042925575.1 |
| ☑ | uncharacterized protein CHLRE_02g143151v5 [Chlamydomonas reinhardtii] | Chlamydomonas reinhardtii | 50.4 | 50.4 | 2% | 2e-05 | 44.23% | 2638 | XP_042927790.1 |

psbS homologs in C. reinhardtii: **psbS1/psbS2** (uniport I.D - A8HPM2, A8HPM5)

**Figure S6. Finding the *C. sorokiniana* CBR and psbS homologs in *C. ohadii* and *C. reinhradtii.* (a)** NCBI protein BLAST analysis of *C. sorokiniana* CBR amino acids sequence vs *C. ohadii* proteome. **(b)** NCBI protein BLAST analysis of CBR homolog of *C. ohadii* vs *C. reinhardtii* proteome. Most similar protein (in black box) was detected in the uniport database, where it was annotated ELIP3. **(c)** NCBI protein BLAST analysis of *C. sorokiniana* psbS amino acids sequence vs *C. ohadii* proteome**.** **(d)** NCBI protein BLAST analysis of psbS homolog of *C. ohadii* vs *C. reinhardtii* proteome**.** Most similar proteins (in black box) were detected in the uniport database, where they were annotated as psbS1 or psbS2

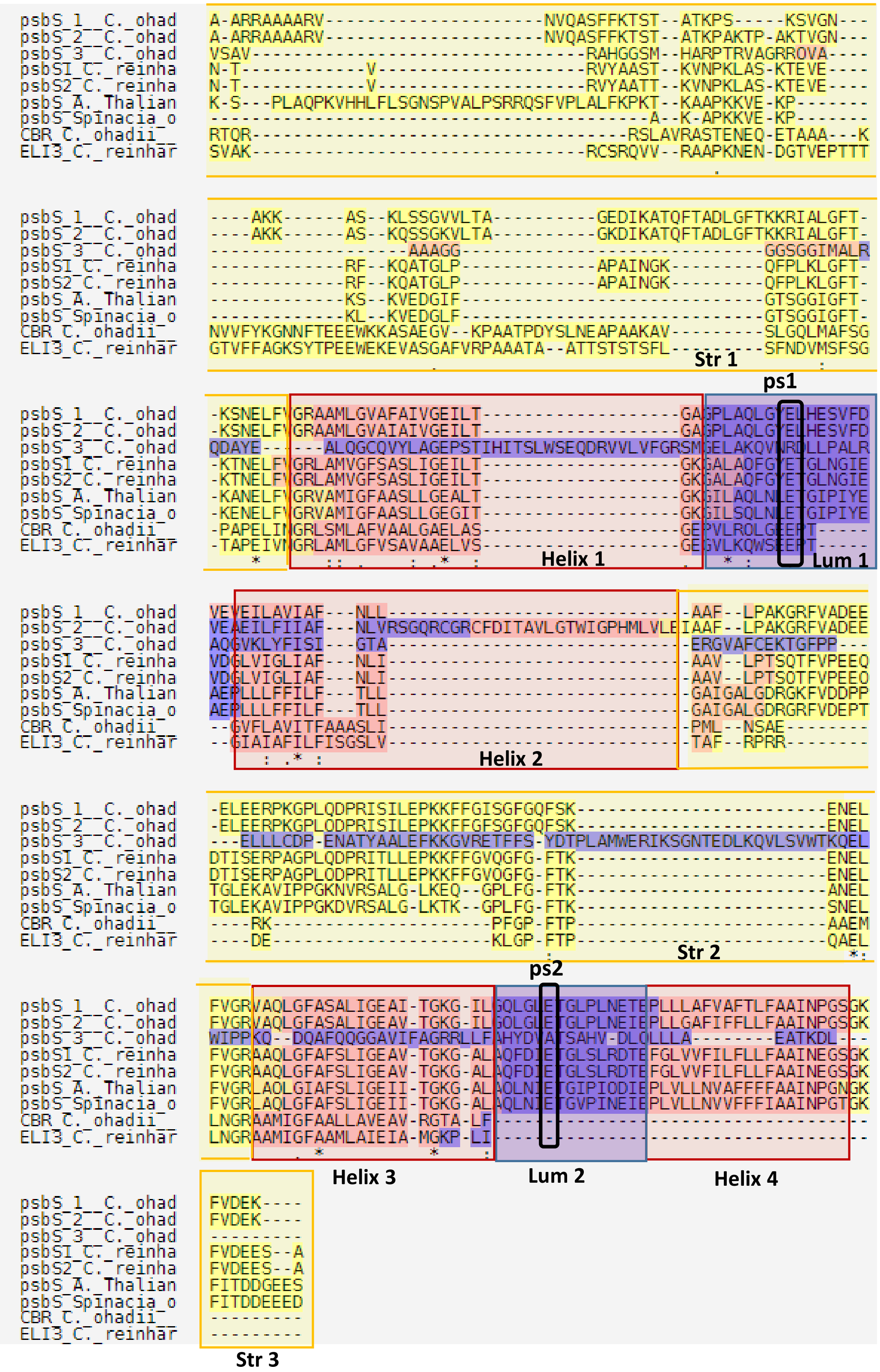


**Figure S7. The *C. ohadii* CBR contains the conserved Glu that functions in proton sensing.** Transmembrane protein multiple sequence alignment of possible homologs to psbS and CBR in *C. ohadii* vs. psbS1, psbs2 and ELIP3 (CBR homolog) of *C. reinhardtii*, psbS of *A. thaliana* and psbS of *S. olerchea.* Note the conserved glutamic acid in CBR (ps1). Str – chloroplast stroma, Lum – Thylakoid lumen, Helix – transmembrane helix, ps – proton sensing.

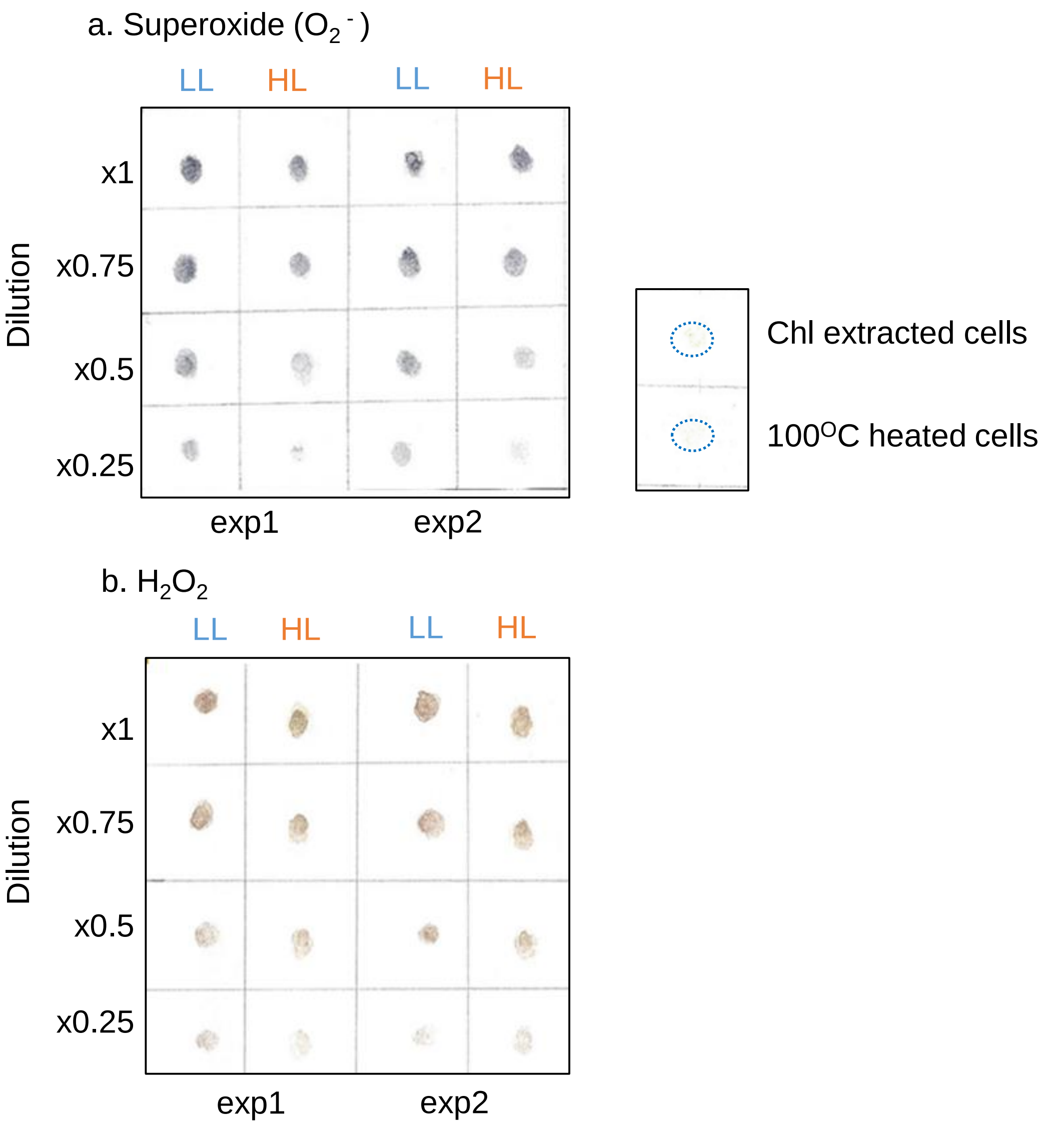


**Figure S8: Accumulation of $O_2^{\cdot -}$ and $H_2O_2$ in LL and HL cells**. **(a)** NBT staining of HL and LL cells illuminated with HL (1000 μmols photons $m^{-2}$ $s^{-1}$) for 10 min. Equal amounts of HL and LL cells were used, which means half of the chlorophyll amount in HL cells as compared to LL cells. Two repetitions of LL and HL experiments, out of four repetitions that were performed, are presented (exp1 and exp2). Each sample was examined in the dilutions of the cells in order to best observe the differences and being able to estimate the percent of difference. Two controls are presented to the right. One in which the chlorophyll was extracted before the experiment and the second which the cells were heated before the experiment. Dashed circles indicate the spots of the cells in the control samples. **(b)** DAB staining of cells as described for panel a. Quantitation of the accumulated NBT and DAB colors is presented in Figure 5.
